# Epigenetic Coalitions Couple Tissue Growth to Generate Periodic Colour Patterns in Birds

**DOI:** 10.64898/2025.12.23.696276

**Authors:** Zhou Yu, Wei Zhao, Chih-Kuan Chen, Wen-Chien Jea, Ya-Chen Liang, Hans I-Chen Harn, Noah Kenneth Brady, Tzu-Yu Liu, Tsz Yau Law, Masafumi Inaba, Ting-Xin Jiang, Ping Wu, Edward Chuong, Qing Nie, Cheng-Ming Chuong

## Abstract

Periodic patterning is fundamental to biological organization. In birds, colour stripes and spots on growing embryonic surface and elongating feather filaments provide a unique system to study continuous periodic patterning in expanding domains. Agouti-signaling protein (ASIP) directly reports epigenomic activity during colour patterning. Using a neural-network-like architecture, we show that ASIP’s *cis*-regulatory landscape functions as hidden layers, integrating morphogen feedback, chromatin topology, and tissue geometry to generate discrete colour outputs. Single-nucleus multiome and Micro-C uncover stage- and context-specific enhancer-silencer coalitions. Epidermal Wnt ligands activate ASIP while inducing Wnt inhibitors in fibroblasts, forming a negative-feedback loop coupling periodic patterning to domain expansion. Comparative cross-tissue and cross-species analyses define *cis*-regulatory modules comprising an epigenetic grammar; functional assays highlight retrotransposon co-option expanding ASIP’s *cis*-regulatory repertoire, potentially contributing to colour pattern evolution. Together, these findings motivate a Turing-principle-based communication model: ASIP reflects epigenetic coalitions shifting to drive diverse, environmentally tunable colour motifs for adaptation.

## Introduction

Tissue patterning is fundamental in both normal development and disease (*1–7*). Among patterning mechanisms, periodic patterning is a major and ubiquitous process that emerges from communication within a cell population. In skin, classic Turing-type morphogen reaction-diffusion networks—often involving Wnt, BMP, FGF and related pathways—can position feather and hair primordia at regular intervals (*8–12*). Similar principles can also be implemented through local cell-cell interactions: in zebrafish, pigment-cell adhesion, repulsion, and signaling—modulated by gap junctions and ion-channel activity—organize stripes and their disruption in mutants (*13*, *14*). In birds, periodic pigment patterns can arise without extensive pigment-cell rearrangements: dermal fibroblasts express Agouti-signaling protein (ASIP), which melanocytes interpret to tune pheomelanin production and generate colour stripes (*6*, *15–18*). In this system, ASIP expression reports the periodic programme in real time, offering a direct view of how epigenomic regulation encodes pattern as tissues grow. Evidence increasingly points to *cis*-regulatory elements (CREs) as key determinants of ASIP expression and pigmentation diversity across species (*17*, *19–25*). More broadly, across development and disease, non-coding enhancers and silencers encode when and where genes are expressed by recruiting transcriptional activators and repressors (*26–32*). This regulatory logic is implemented within three-dimensional (3D) genome architecture, which organizes loci into interaction networks that enable long-range CRE-promoter communication and precise spatiotemporal control of transcription (*33–41*).

Beyond these specific mechanisms, developing tissues do not simply execute a genetic script; they compute form. Cells continuously integrate biochemical and biophysical cues, exchanging information across space and time to assemble coherent patterns (*42*). From this perspective, periodic patterns reflect multiscale coordination: they emerge when morphogen cues, growth dynamics, and chromatin regulation are coupled in a self-organizing circuit, rather than a single linear cascade (*42–44*).

Avian skin provides a powerful system to dissect periodic patterning logic for two key reasons. First, ASIP serves as a direct and spatially periodic readout of a Turing-like system. Turing principles have been invoked for hair, feather, and intestinal villus patterning, but typically as upstream of multiple intervening processes such as cell migration and bud morphogenesis. In contrast, ASIP expression in avian skin directly reports the underlying epigenomic activity, enabling interrogation of Turing-like logic at the epigenetic *cis*-regulatory and chromatin levels. Second, this patterning occurs within a growing tissue. Classical Turing models generally assume a static field, whereas skin expands continuously during development (*45*). Birds provide two natural “growing domains”: ASIP stripes emerge sequentially across the two-dimensional embryonic dorsum to form a macro-scale body pattern, while discrete dots appear along the one-dimensional growing feather filament to generate a micro-scale within-feather pattern (*6*, *15*, *16*, *46–49*). Yet, despite these systems enabling explicit links between domain growth, inhibitor feedback, and stripe insertion, it remains unknown how such tissue-scale signals are integrated at a single genomic locus to produce temporospatial ordered ASIP expression domains.

To address this question, we integrate single-nucleus multiome profiling across embryonic stages with Micro-C mapping of chromatin topology, functional dissection of CREs, transcription factor perturbations, comparative analyses across tissues and species, and mathematical modeling. These approaches connect epigenomic dynamics at cell-state resolution to 3D genome organization and regulatory function. They reveal two epigenetic layers of ASIP regulation: CRE coalitions—context-dependent assemblies of enhancers, silencers, and dual-function elements—and CRE modules, sets of elements whose accessibility shifts in concert across contexts. We show that the ASIP locus functions as a multilayer integrator, implementing a reaction-diffusion programme, in which dynamic enhancer-silencer coalitions couple morphogen feedback to tissue geometry and chromatin topology. Together, these findings suggest that multimodal *cis*-regulatory coalitions offer a flexible, evolvable framework for encoding periodic gene expression in growing tissues—one that supports both the robustness and diversity of colour patterns across species.

## Results

### 1. Single-nucleus multiome profiling maps ASIP CREs in embryonic quail skin

To define the regulatory architecture underlying periodic ASIP expression, we first established a developmental and spatial framework for interpreting *cis*-regulatory activity. In the Japanese quail embryo, ASIP expression follows a reproducible temporal sequence that provides such a reference (Fig. 1A). There are longitudinal stripes across the dorsal skin in land fowl (Fig. 1B) (*6*, *16*). In Japanese quail, we will focus on the three sets of stripes that appear in the dorsal feather tract sequentially. The first pair of ASIP stripes (A1) emerges bilaterally along the lateral flanks at embryonic day 4 (E4). This is followed by a midline stripe (A2) at E5. Then a third set (A3) arising between A1 and A2 around E5.5 as the skin field expands (Fig. 1, A and C). A distinct stripe appears on the leg (AL) appear in the femoral tract (Fig. 1C). These developmental landmarks define a coordinate system on the body surface for mapping regulatory dynamics as periodic stripes are progressively established.

**Fig. 1.**
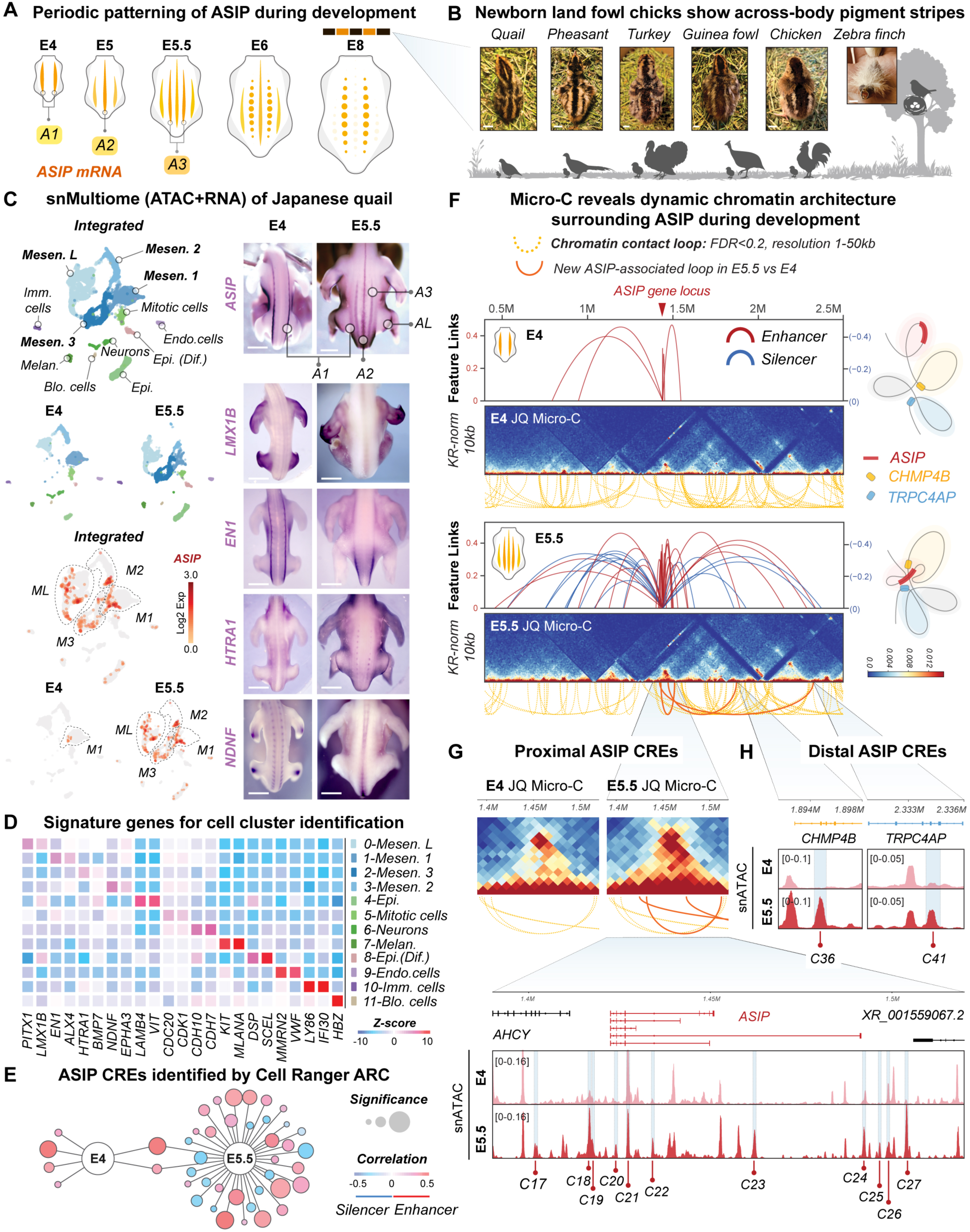
snMultiome identifies CREs underlying periodic ASIP expression in embryonic quail skin. **A,** Schematic illustrating temporal-spatial patterning of ASIP expression across the expanding embryonic quail skin domain. **B**, Newly hatched land fowl chicks display across-body pigment stripes **C,** snMultiome (ATAC + RNA) analysis of Japanese quail skin at E4 and E5.5, showing WNN-UMAP projections integrating snATAC-seq and snRNA-seq data. Whole-mount RNA *in situ* hybridization for ASIP, LMX1B, EN1, HTRA1, and NDNF identifies distinct mesenchymal populations. **D,** Heatmap of snMultiome data showing cluster-specific marker gene expression. **E,** Developmental-stage-dependent ASIP CREs identified using Cell Ranger ARC. **F,** Micro-C analysis reveals dynamic chromatin architecture surrounding the ASIP locus during development. **G,** Distribution of proximal ASIP CREs near the ASIP locus. **H,** Distal ASIP CREs exhibiting long-range chromosomal contacts with ASIP, revealed by Micro-C. Credit: B, photos of day-old pheasant, turkey, and zebra finch from Yu (*108*). C, ASIP *in situ* modified with permission from Inaba et al (*16*). Scale bars: 1 cm (B), 2 mm (C).

To map cell states and their regulatory landscapes, we applied single-nucleus multiome profiling (snRNA-seq + snATAC-seq) and integrated modalities with weighted nearest-neighbor (WNN) analysis (Fig. 1C) (*49*). WNN jointly learns relationships between transcriptomic and chromatin-accessibility profiles, allowing cell clustering to reflect shared regulatory and transcriptional identities rather than either modality alone. We sampled two representative timepoints, E4 and E5.5, capturing the onset of A1 and the transitional phase when A1 diminishes as A2 and A3 consolidate in the expanded skin field (Fig. 1C) (*16*). Joint clustering revealed four mesenchymal populations whose spatial distributions align with the four ASIP stripes observed *in vivo* (Fig. 1C and fig. S1A). We then mapped these WNN-defined mesenchymal cell clusters to anatomical positions by RNA *in situ* hybridization of cluster-enriched markers: EN1 marked M1, NDNF marked M2, HTRA1 marked M3, and LMX1B marked the leg-associated cluster ML (Fig. 1, C and D, and fig. S1B). Because each cluster occupies a characteristic position relative to ASIP expression, we name the clusters according to the ASIP stripe domain they contain. At E4, only A1 is present and corresponds to M1. By E5.5, two additional ASIP domains (A2 and A3) appear medial to A1 and map to M2 and M3, respectively, and the distinct leg stripe (AL) corresponds to ML (Fig. 1C). Notably, EN1 not only delineates the M1 compartment but also exhibits a stripe-like pattern that parallels ASIP at E4 and diminishes as A1 faded by E5.5, implicating EN1 as an upstream transcriptional activator of ASIP (Fig. 1C).

Because CREs govern where and when genes are expressed, identifying the CREs that control ASIP is essential for understanding how periodic patterns are encoded. In this context, we use “enhancers” to refer to *cis*-regulatory elements that increase transcription probability, and “silencers” to refer to elements that repress or restrict gene expression; both are defined by chromatin accessibility and their correlation with gene output (*50*, *51*). We further define a “CRE coalition” as a coordinated set of CREs—a mixture of enhancers and silencers—whose combined epigenetic activity shapes a specific pattern output. Our working hypothesis is that the emergence and refinement of periodic ASIP stripes arise from the sequential engagement of specific enhancers and silencers during development. To test this hypothesis, we next asked how CREs near ASIP become accessible (open) and function across developmental time. Using Cell Ranger ARC (*49*, *52*), we performed peak-to-gene linkage analysis to correlate chromatin-accessibility peaks with ASIP expression across single nuclei, nominating CREs whose accessibility dynamics are most likely to influence ASIP transcription. This analysis identified 42 candidate ASIP CREs across E4 and E5.5 stages, designated C1 through C42 (Fig. 1E). Two predicted enhancers were shared between E4 and E5.5, whereas stage-specific repertoires dominated. At E4, we detected 6 predicted enhancers and notably no silencers, consistent with the hypothesis that early ASIP activation occurs within an enhancer-permissive chromatin landscape. By E5.5, the repertoire expanded to 24 predicted enhancers and 14 predicted silencers, revealing a developmentally delayed opening of silencer elements relative to enhancers. One can see the large expansion of ASIP CREs from E4 to E5.5 (Fig. 1E), The temporal separation between enhancer and silencer activation also suggests distinct regulatory phases in which ASIP is first broadly potentiated and then progressively refined as the pattern proceeds.

To relate three-dimensional genome architecture to these regulatory changes, we performed Micro-C on matched E4 and E5.5 skin mesenchyme samples. Between E4 and E5.5, the topologically associating domain (TAD) encompassing ASIP exhibited largely increased contact frequency within the domain and strengthened interactions between ASIP and nearby accessible CREs (Fig. 1, F to H). At E5.5, these new loop anchors emerged at the ASIP locus, generating E5.5-specific loops that were not detectable at E4. In addition to these local structural changes, Micro-C revealed two long-range interactions that defined distal regulatory neighborhoods: C36 near CHMP4B and C41 near TRPC4AP (Fig. 1H). Together, these data indicate that the ASIP locus is incorporated into a progressively more interconnected regulatory architecture during development (Fig. 1G, bottom panel, E5.5 track), providing both local and distal contact scaffolds through which enhancers and silencers can exert stage-specific effects. Integrating with accessibility data, the 3D contact landscape suggests that ASIP expression is initially facilitated by enhancer accessibility and subsequently refined by silencer engagement and the maturation of chromatin topology.

### 2. CRE coalitions and Wnt signaling jointly regulate ASIP patterning in a growing domain

Figure 1 shows that ASIP is regulated by a diverse, stage-dependent repertoire of CREs that are organize into dynamic coalitions with distinct accessibility profiles over developmental time. This complexity raised the possibility that ASIP regulation is not governed by isolated elements acting independently, but instead by combinations of CREs that operate cooperatively or antagonistically within each stripe-forming mesenchymal population. This raised two central questions: how does each stripe assemble its specific CREs coalition, and what regulatory principles are shared across stripes? Because peak-to-gene linkage identifies CREs correlated with ASIP across all mesenchymal cells, we reasoned that a stripe-specific analysis was needed to resolve which CREs are actively deployed within each stripe identity. To address this, we applied DIRECT-NET (*53*), a population-specific CRE discovery pipeline that identifies and categorizes CRE sets assembled by ASIP-expressing cells in each mesenchymal population.

We refer to these population-defined sets as stripe-specific CRE coalitions—groups of *cis*-regulatory elements that act together to drive stripe-specific ASIP expression. Across M1, M2, M3, and ML, DIRECT-NET identified 73 non-redundant ASIP CREs (Fig. 2A). Of these, 26 (35.6%) were unique to a single population, whereas the remainder were shared to varying degrees: 26 (35.6%) shared by two groups, 11 (15.1%) by three, and 10 (13.7%) by all four (Fig. 2B). Functional classification based on accessibility-expression correlations further revealed that among the 47 shared CREs, 21.3% behaved as silencers, 8.5% as enhancers, and 70.2% exhibited dual-function behavior—capable of acting as either enhancer or silencer depending on mesenchymal context (Fig. 2B). These results indicate that functional plasticity of CREs is a major feature that enables dynamic periodic expression of ASIP.

**Fig. 2.**
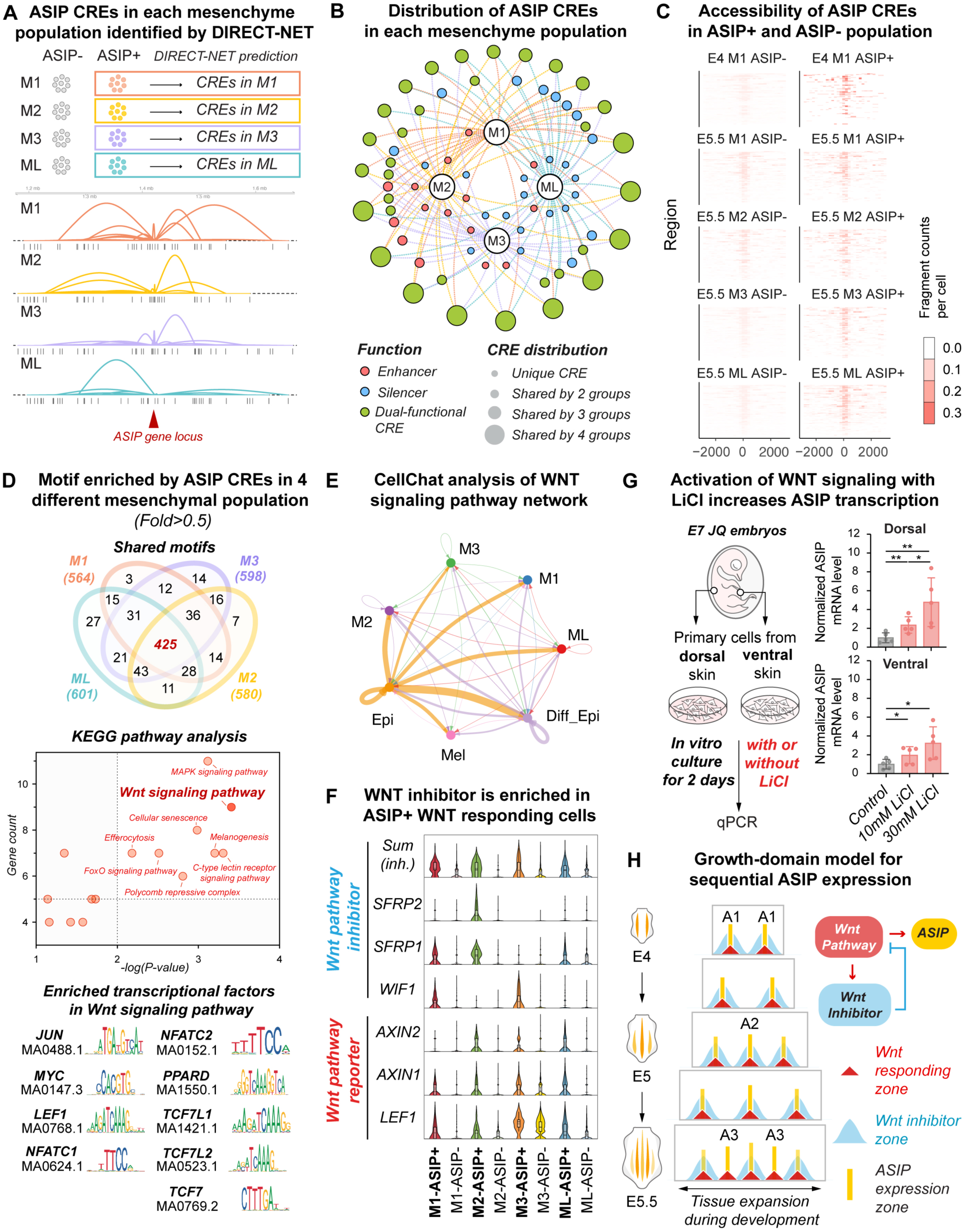
ASIP CRE coalitions across mesenchymal populations respond epidermal derived Wnt signaling to generate sequential ASIP expression in the expanding skin domain. **A,** ASIP CREs from temporo-spatial ASIP stripes are identified in each mesenchymal cell clusters by DIRECT-NET. See Fig. 1A and 1C for the spatiotemporal emergence of mesenchymal stripes (M1–3, L) containing corresponding ASIP stripes (A1–3, L). **B,** Distribution and functional annotation of the above ASIP CREs across mesenchymal populations. **C,** Accessibility of ASIP CREs in ASIP+ versus ASIP-cell clusters in different colour stripes. **D,** Motif-enrichment analysis of ASIP CREs highlights Wnt signaling as a shared regulatory axis across four mesenchymal populations underlying yellow stripes. **E,** CellChat analysis shows that Wnt signaling is a key regulator of ASIP-expressing mesenchymal cells. **F,** Violin plots show expression of Wnt inhibitors and Wnt reporters in ASIP+ and ASIP-cells across populations. **G,** Activation of Wnt signaling with LiCl increases ASIP transcription (mean ± s.d., n = 5, two-tailed t-test). **H,** Growth-domain model for sequential ASIP expression.

We then asked whether chromatin accessibility itself operates as a strict on-and-off switch for ASIP. Across mesenchymal populations, most coalition CREs remained accessible in both ASIP+ and ASIP− cells, though at consistently higher accessibility in ASIP+ cells (Fig. 2C and fig. S2). This persistent accessibility suggests a permissive chromatin state that supports activator-repressor competition (*51*, *54*, *55*) and enables rapid shifts in ASIP output as signaling inputs continue to change with expansion of tissue domain.

To further identify regulatory logic shared across stripe identities, we analyzed transcription factor motifs enriched within each stripe-specific CRE coalition. Motifs enriched by more than 0.5-fold were compared across populations (Fig. 2D). KEGG analysis of motifs shared by all four populations pointed to Wnt-associated transcription factors, including JUN, MYC, LEF1, NFATC1, NFATC2, PPARD, TCF7L1, TCF7L2, and TCF7 (Fig. 2D, lower panel). Consistent with this, communication analysis using CellChat (*56*) identified epidermal cells as the major source of Wnt ligands for the mesenchymal cell populations (Fig. 2E and fig. S3A), in alignment with the epidermis-dermis dialogue that patterns developing skin. Within each mesenchymal group, Wnt reporter genes (Axin1, Axin2, Lef1) were found higher in ASIP+ than ASIP− cells (Fig. 2F). Intriguingly, secreted Wnt inhibitors were also elevated in ASIP+ cells, but with stripe-specific combinations: M1 expressed Wif1 and Sfrp1, M2 expressed Sfrp1 and Sfrp2, M3 expressed Wif1, and ML expressed Sfrp1(Fig. 2F and fig. S3B). Summed inhibitor levels were consistently higher in ASIP+ than in ASIP− cells within the same population, pointing to a negative feedback architecture in which Wnt-responsive mesenchyme locally produces antagonists to define and refine ASIP domains.

To test whether Wnt activity is upstream of ASIP transcription, we activated Wnt signaling in primary Japanese quail E7 dorsal and ventral skin cultures using LiCl, a GSK3β inhibitor (*57–60*). LiCl increased ASIP expression in primary cells from both regions in a dose-dependent manner (Fig. 2G). Although ventral-derived cells had higher baseline ASIP, LiCl further increased ASIP expression (fig. S3C). These findings indicate that Wnt activation increases ASIP transcription in both dorsal and ventral skin, regardless of their different baseline expression levels.

Together, these findings support a growth-domain model in which epidermal Wnt input, mesenchymal Wnt sensing, and local feedback inhibition converge on ASIP CRE coalitions to stabilize and position ASIP stripes as the tissue expands (Fig. 2H). Early ASIP+ mesenchyme (A1) is Wnt-responsive and simultaneously secretes Wnt antagonists, establishing flanking inhibitory zones. As the domain enlarges, dilution of inhibitors at the midline enables a permissive corridor for A2 activation. Continued growth and inhibitor turnover then produce additional permissive zones between A1 and A2, enabling A3 formation. In this framework, stripe timing and position emerge from moving reaction-diffusion thresholds, encoded in a CRE-based epigenetic substrate and tuned by inhibitor dynamics that change as the tissue grows.

### 3. A dual-colour ratiometric reporter dissects enhancer-silencer dynamics at ASIP CREs

Although coalition-level analyses reveal how mesenchymal cell populations deploy distinct combinations of CREs, they do not, by themselves, resolve the intrinsic regulatory logic of individual elements. Correlations in accessibility cannot distinguish whether a CRE is activating, repressing, or context-dependent, nor can they reveal competition between enhancer and silencer activities that may be embedded within the same sequence. To define these element-level rules, we combined functional reporter assays and transcription-factor perturbations to quantify enhancer-silencer behavior at single-element resolution, and subsequently applied these rules to understand how individual elements logic scales to coalition-level patterning.

To accomplish this, we engineered a dual-colour ratiometric reporter capable of quantitatively dissecting intrinsic enhancer-silencer logic at single elements (Fig. 3A and fig. S4). Most candidate CREs in our dataset are ∼800 bp, consistent with the typical size of vertebrate enhancers (*50*, *61*, *62*). Candidate CREs were placed upstream of a miniCMV promoter driving the short half-life d2EGFP, enabling sensitive detection of both activation and repression on a weak basal background (*63*, *64*). A parallel mCherry cassette driven by the CAG promoter served as an internal normalization control. The two cassettes were separated by the chicken β-globin 5’HS4 insulator to minimize regulatory crosstalk, and the entire reporter was flanked by additional β-globin 5’HS4 insulators to buffer genomic position effects upon Tol2-mediated integration (*65–67*). Reporter activity was quantified as the d2EGFP/mCherry ratio, normalized to a non-CRE control (Fig. 3A).

**Fig. 3.**
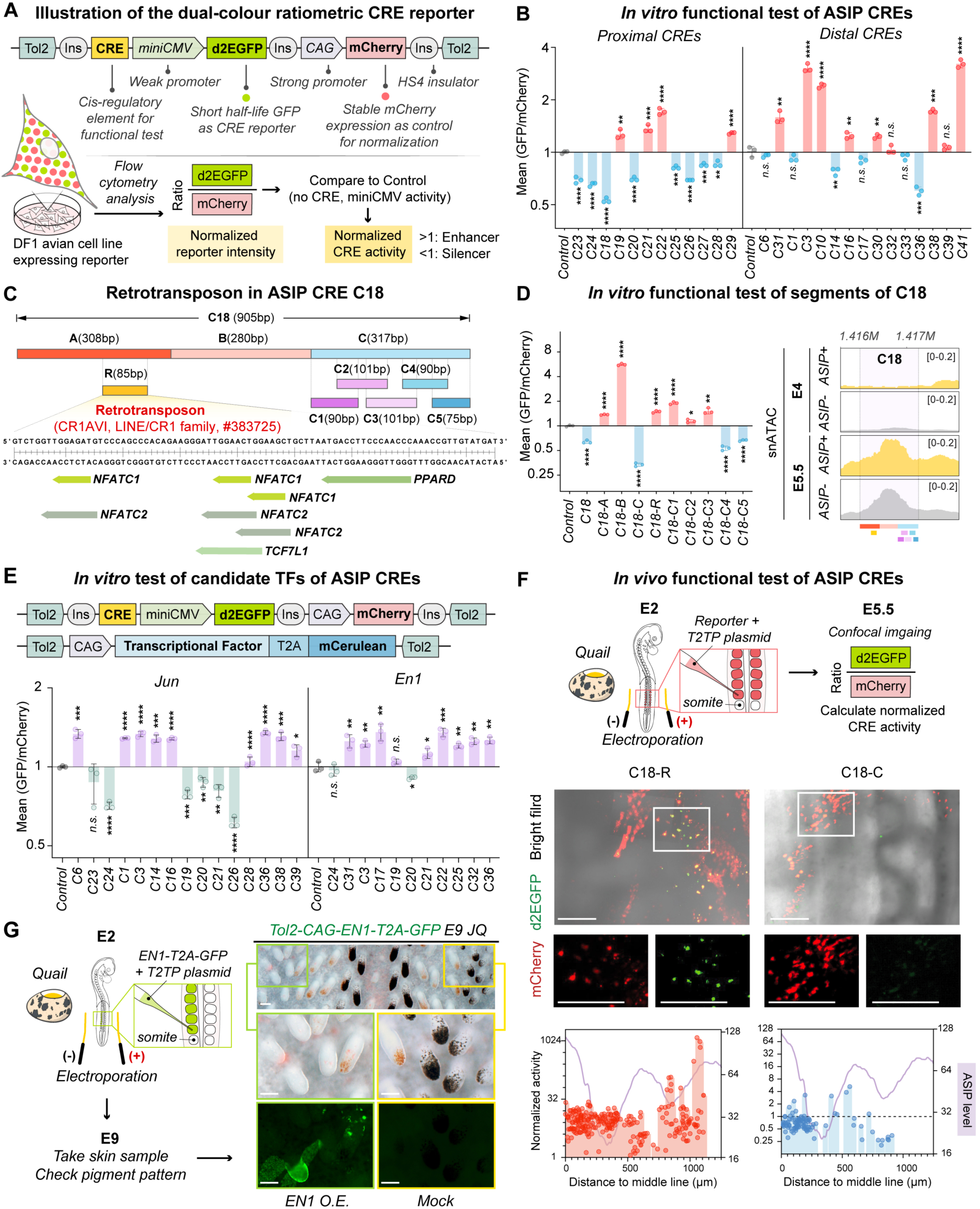
Functional characterization of ASIP CREs using a dual-colour ratiometric reporter system. **A,** Schematic of the dual-colour ratiometric CRE reporter design and experimental workflow for assessing CRE activity. **B,** *In vitro* functional results of ASIP CREs in DF-1 avian cells. **C,** Retrotransposon #383725 (LINE/CR1 family) is a major component of ASIP CRE C18. **D,** Functions of C18 CRE fragments are analyzed *in vitro* using DF-1 cells. **E,** Function of candidate transcription factors (e.g., Jun and En1) on CRE activity are analyzed using the ratiometric reporter system. **F,** *In vivo* functional assay of ASIP CREs via somite injection and electroporation in quail embryos. C18-R functions as an enhancer, whereas C18-C acts as a silencer. **G,** Overexpression of EN1 alters pigment patterning. Data are mean ± s.d. (B, D, E); n = 3; two-tailed t-test; > 1000 cells analyzed per replicate by flow cytometry. Scale bars: 100 µm (F, G).

We apply this system to DF-1 avian fibroblast cells and found three unexpected features. First, many elements originally annotated as enhancers functioned as silencers (Fig. 3B, CREs in blue). This shift suggested that the regulatory identity of a CRE cannot be inferred from peak accessibility alone and may reflect the combined effects of multiple embedded regulatory components (*26*, *68*). Given the dual-function properties observed at the coalition level, we reasoned that individual CREs might harbor both activating and repressing components whose combined output determines the summed activity of the full-length element. To test this possibility, we dissected representative CREs into smaller fragments to map internal subregions and identify enhancer- and silencer-bearing components, thereby revealing how their interactions generate the composite regulatory behavior of each CRE. For example, dissecting the strongest silencer, C18, showed that ∼300-bp fragments C18A and C18B behaved as enhancers, whereas C18C carried silencing activity, with repression mapped by tiling to C18C4-C18C5 at the 3′ end (Fig. 3, C and D).

Second, CRE behavior was non-additive: the activity of full-length C18 could not be predicted from the activities of its constituent parts (Fig. 3, C and D), indicating that enhancer and silencer components within the element influence each other’s output.

Third, we identified a retrotransposon-derived enhancer within C18. C18R, an 85-bp CR1AV1 (LINE/CR1) cassette within C18A, was sufficient to boost reporter expression. Motif scanning with nine Wnt-associated transcription factors (Fig. 2D) revealed that NFATC1, NFATC2, TCF7L1, and PPARD motifs embedded within this 85-bp cassette (Figs. 3C). Comparative analysis showed that this element is conserved across land fowls—including quail, chicken, guinea fowl, and turkey (fig. S5). Its most conserved core (GATTGGAACTGGAAGCT) contains motifs for NFATC1, NFATC2, and TCF7L1, suggesting a conserved *cis*-regulatory logic across species.

Consistent with this competition model, snATAC showed that at E4, there is no accessibility at C18 in either ASIP+ or ASIP− cells, but equal opening in both cell types at E5.5 (Fig. 3D, right panel). Thus, failure to express ASIP cannot be attributed solely to CRE closure; rather, the balance of activators and repressors on an open scaffold determines output. We therefore tested the impact of candidate transcription factors (TFs) on CRE activity by co-expressing each TF with the CRE reporters. To this end, we used a second Tol2 construct that co-expressed the transcription factor with mCerulean via a T2A peptide. We examined EN1, associated with the A1 stripe, and JUN, a Wnt-pathway effector (Fig. 3E and fig. S4B). In DF-1 cells, EN1 acted predominantly as an activator across tested CREs, whereas JUN showed bidirectional effects, activating some CREs and repressing others to comparable degrees (Fig. 3E). *In vivo*, somite injection and electroporation to overexpress EN1 produced yellow to pale patches on the manipulated side (Fig. 3G, right panel, left photo column) relative to the contralateral control, consistent with ASIP overexpression (*16*).

We further tested ASIP CRE activity *in vivo* as it offers key advantages: it enables monitoring of CRE function in primary cells within their native developmental niche and preserves spatial context, allowing CRE activity to be mapped to ASIP transcriptional output measured by *in situ* hybridization (Fig. 1C). The reporter plasmids were delivered into E2 Japanese quail somite by electroporation, which subsequently give rise to mesenchymal cells (Fig. 3F, upper panel). Embryos were then collected and live-imaged at E5.5 by confocal microscopy to quantify reporter fluorescence. As in the *in vitro* quantification assay, normalized CRE activity >1 was classified as enhancer activity, whereas activity <1 was classified as silencer activity (Fig. 3, A and F).

In E5.5 quail skin, C18R and C18C reporters corroborated the *in vitro* results we observed in chicken cells (Fig. 3D), supporting conservation of CRE function across species (Fig. 3F, lower panel). C18R showed normalized activity above 1 across reporter-expressing cells, indicating that the retrotransposon functions as an enhancer. In contrast, C18C exhibited normalized activity below 1 in most reporter-expressing cells, consistent with its silencer activity *in vitro*. Notably, when CRE activity was analyzed together with spatial position (distance to the midline) and local ASIP transcriptional levels (measured from ASIP *in situ* hybridization), activity of a single CRE was not always concordant with ASIP expression, despite its overall enhancer or silencer function (Fig. 3F, lower panel). These results further support that a multi-CRE coalition, rather than any single element, determines the final ASIP transcriptional output.

Together, the reporter and TF-perturbation experiments demonstrate that individual ASIP CREs comprise interacting enhancer and silencer component whose combined output is context-dependent and non-additive. This intrinsic modularity provides a mechanistic basis for the dual-function behavior observed at the coalition level and explains how the same locus can generate distinct transcriptional states across a growing tissue. By demonstrating that CREs operate as competitive integration hubs—rather than as simple enhancers or silencers, these findings establish a core principle governing ASIP’s dynamic expression and offers a mechanistic framework for how reaction-diffusion logic is executed at epigenomic level to produce periodic patterns.

### 4. Cross-species multiome profiling reveals evolutionary co-option of transposons-derived ASIP enhancers

Avian periodic colour patterns can be organized at multiple scales, from across-the-body striping to micro-scale motifs within a single feather vane (*45*). To determine whether the principles uncovered from embryonic pigment stripes are likewise employed in within-feather patterning, we performed snMultiome profiling of regenerating adult guinea fowl feathers five weeks after plucking (Fig. 4, A to B, and fig. S6). Guinea fowl feathers exhibits a micro-scale polka-dot pattern within each feather (Fig. 4A). We then asked how ASIP are regulated in this morphogenetic context.

**Fig. 4.**
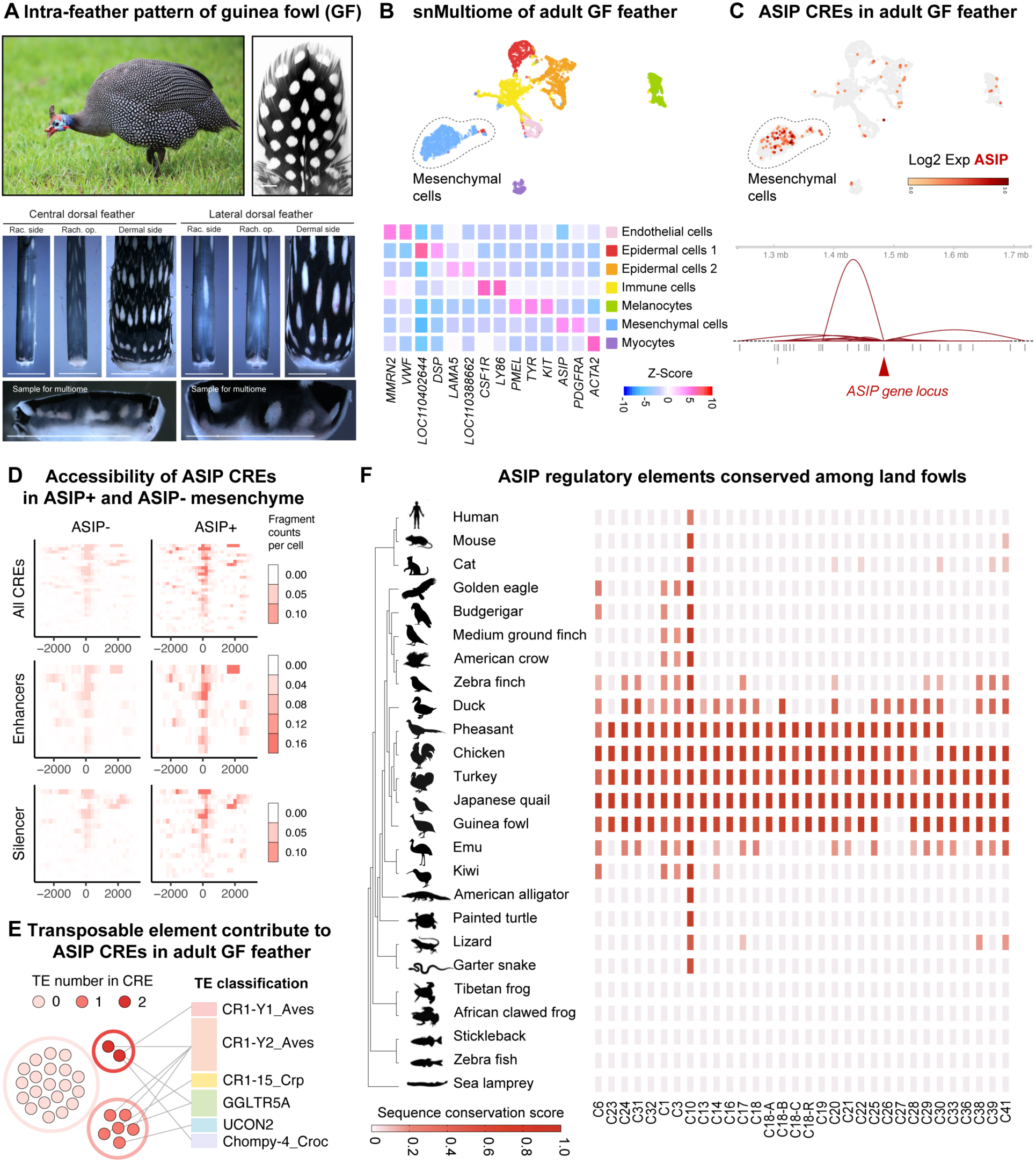
snMultiome analysis of adult guinea-fowl feathers reveals evolutionary adoption of transposon elements in ASIP CREs for intra-feather patterning. **A,** A wild guinea fowl (Swakopmund, Namibia) exhibiting periodic white-dot pigment patterns on feathers. **B,** snMultiome analysis of adult guinea fowl feathers, with heatmap showing signature genes for cell-cluster identification. **C,** ASIP-expressing cell clusters in feather pulp mesenchyme and corresponding ASIP CREs are identified by DIRECT-NET. **D,** CRE accessibility in ASIP+ and ASIP-feather pulp mesenchymal cells. **E,** Different levels of transposon contribution to ASIP CREs in adult guinea fowl feathers. **F,** Conservation of ASIP CREs among land fowl species exhibiting transverse pigment stripes. Credit: A, guinea fowl photo by Yannick Francois / Macaulay Library, Cornell Lab of Ornithology (ML616821372). Scale bars: 5 mm (A).

Using DIRECT-NET, we identified 28 ASIP CREs in guinea fowl feather mesenchyme (Fig. 4C, lower panel). Of these, 9 (32.1%) predicted primarily as enhancers and 19 (67.9%) as silencers (Fig. 4D). Interestingly, 27 of the 28 elements exhibited sequence conservation in Japanese quail (Fig. 5B), and three overlapped with quail CREs identified in embryonic skin (Fig. 5B). Thus, the ASIP CRE coalition operating in regenerating guinea fowl feathers mirrors the ASIP CRE coalition in the embryonic Japanese quail skin. These results imply that a partially shared regulatory toolkit is deployed across species and across distinct ASIP patterning contexts.

**Fig. 5.**
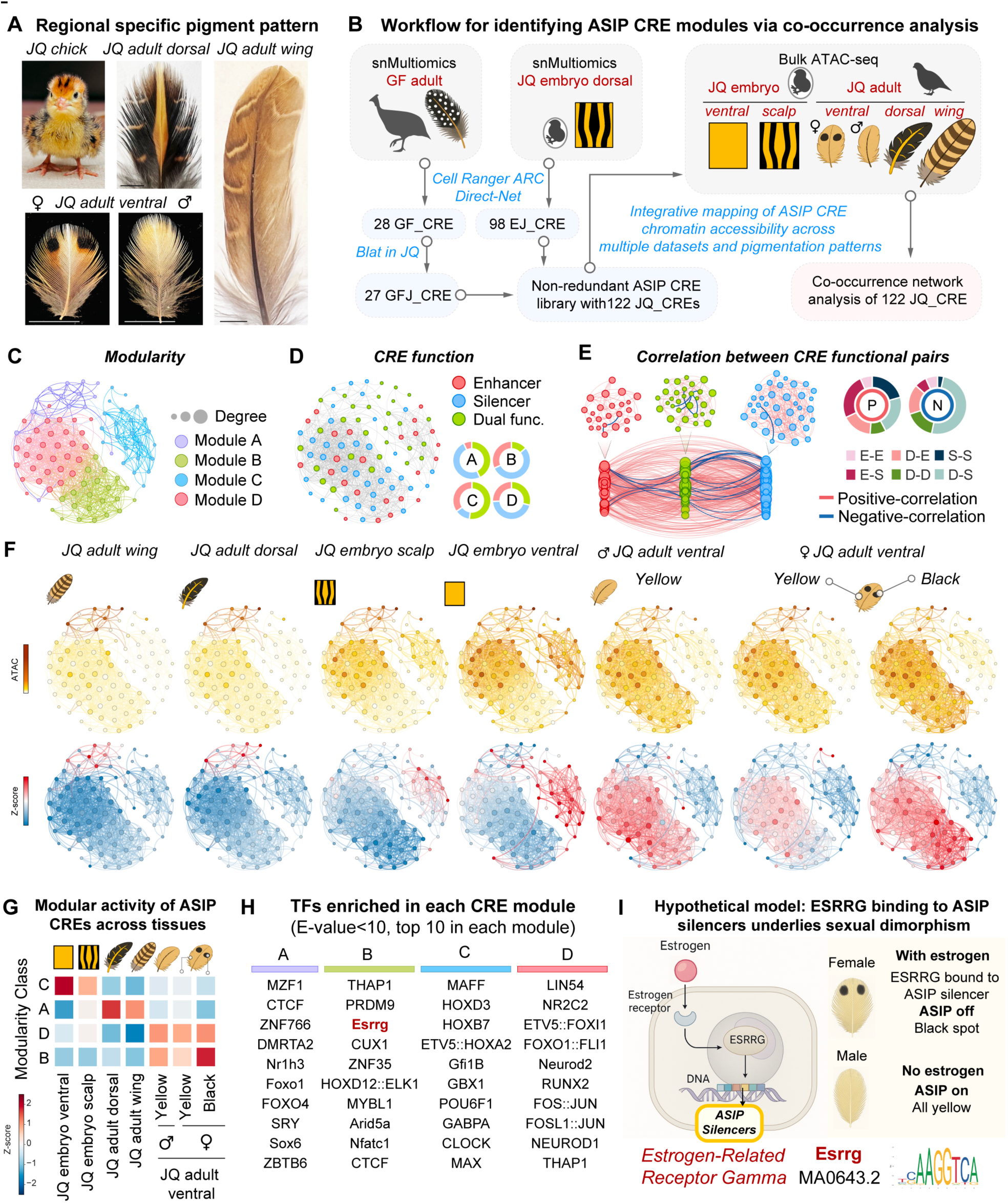
A spectrum of periodic pigment patterns across skin regions, developmental stages, and land fowl species arises from ASIP CRE interactions within multimodal regulatory coalitions. **A,** Regional specific and sexual dimorphic pigment patterns of Japanese quail. **B,** Workflow for identifying ASIP CRE modules via co-occurrence analysis. **C,** Correlation-based clustering segregates ASIP CREs into four CRE modules (A-D). **D,** Functional classification of CREs as enhancers (red nodes), silencers (blue nodes), or dual-function elements (green nodes). Donut plots depict the relative proportions of each category within modules. **E,** CRE correlations classified as positive (red line) or negative (blue line), with connection types quantified in donut plots. Nodes denote enhancers (red), silencers (blue), and dual-function elements (green). **F,** Distinct spot and stripe colour patterns and their distinct communication patterns across ASIP CRE modules. ATAC tracks display normalized ATAC-seq reads per CRE; Z-scores were calculated per CRE across 7 tissues. **G,** Modular activity of ASIP CREs across tissues. Average occupancy of all CREs within each module is shown; Z-scores calculated across 7 tissues. **H,** Transcription factors enriched in different CRE modules. **I,** Hypothetical model illustrating black-dot formation in female adult ventral feathers. Abbreviations: P, positive correlation; N, negative correlation; E, enhancer; S, silencer; D, dual-functional. Scale bars: 5 mm (A).

One striking feature we found about the guinea fowl repertoire is its transposon content: six CREs harbor one transposable element each, and two harbor two elements (Fig. 4E and table S1). Together with the retrotransposon-derived enhancer module in quail C18R, these findings suggest that transposon-derived sequences are recurrently co-opted as regulatory modules in ASIP CRE evolution, supplying latent transcription factor motifs that can be incorporated to enrich CRE coalition repertoire for pattern generation. Along this line, sequence conservation analysis using BLAT (*69*) indicated that ASIP CREs are broadly conserved among land fowl species that exhibit conserved transverse striping during embryonic development (Fig. 4F). Thus, Fig. 4 shows that ASIP regulation is built on a conserved *cis*-regulatory scaffold and incorporates evolvable components—including transposon-derived elements—that can be repurposed across developmental contexts and species to generate diverse periodic motifs ranging from stripes to dots.

### 5. Cross-tissue network analysis identifies ASIP CRE modules underlying pigment pattern diversity

Japanese quail exhibit striking pattern diversity across development and body regions: hatchlings display yellow ventral downy feathers; dorsal and scalp regions display feathers with transverse stripes; adults acquire thin yellow pinstripes on dorsal and wing feathers; and adult females, not males, develop two black dots on ventral feather (Fig. 5A). While Figure 1–4 identified CRE coalitions of ASIP regulation in embryonic stripes and its evolutionary flexibility across species, we wonder how a single ASIP locus generates this full spectrum of pigment architectures across tissues, developmental stages, and sexes.

To address this, we required a unified view of all ASIP regulatory elements so that CRE deployment could be compared across patterning contexts. We therefore assembled an integrated ASIP CRE library by merging embryonic quail CREs with adult guinea fowl feather CREs, removing redundancies to yield 122 non-overlapping CREs (Fig. 5B). This unified library provides a shared regulatory vocabulary for cross-tissue analyses. We next asked how different pigment-producing tissues deploy distinct subsets of these CREs. Because CREs can only act when accessible, we profiled mesenchymal chromatin accessibility by bulk ATAC-seq across seven major pigment contexts—E7 ventral skin, E7 scalp skin, adult dorsal feather, adult wing flight feather, adult male ventral feather, and adult female ventral feather separated into yellow and black regions (Fig. 5B). Together, these samples capture a wide range of ASIP-dependent pigment architectures in quail.

We quantified accessibility of 122 ASIP CREs across 7 tissues and applied co-occurrence network analysis (*70*, *71*) to identify groups of CREs with correlated accessibility across tissues. Here we define these groups as “CRE modules”: sets of CREs whose accessibility changes in concert across contexts—opening and closing together, or in opposite directions—suggesting shared upstream inputs within modules and distinct upstream inputs across modules. Using this approach, we identified four ASIP CRE modules, A-D (Fig. 5C). Four observations outline the interaction structure of these ASIP modules. First, each module contains a mixture of enhancers, silencers, and dual-function elements (Fig. 5D). Most CRE pairs show positive correlations in accessibility, whereas a smaller subset shows negative correlations, suggesting that some elements may compete for accessibility or operate in a mutually exclusive manner (Fig. 5E). Second, all six types of CRE pairings—including enhancer-enhancer, silencer-silencer, enhancer-silencer, and pairings involving dual-function elements—occur across both positive and negative edges, indicating that CRE relationships span functional classes and likely depend on context (Fig. 5E). Third, negative correlations are observed not only between enhancers and silencers but also within each class, which is consistent with potential competition for shared cofactors or signaling inputs (Fig. 5E). Fourth, dual-function elements frequently connect multiple modules, raising the possibility that individual sequences may adopt different roles across developmental stages, tissues, or signaling environments (Fig. 5, D and E); such context-dependent regulatory switches could offer a parsimonious route to a wide palette of periodic motifs from a compact toolkit. Collectively, these observations support a view of a distributed interaction landscape in which enhancers and silencers engage in both cooperative and competitive ways, enabling more complex regulatory interactions to shape ASIP expression patterns.

Visualization of z-scored accessibility showed that the four modules are differentially deployed across a broad spectrum of patterning contexts (Fig. 5, F and G). Notably, the recurrence of the same module across contexts suggests that they are reused and recombined to build distinct patterns. We therefore asked which upstream transcription factors provide module-specific inputs, and performed TF-motif enrichment analysis for each module to nominate candidate regulators. For example, in Module B (most accessible in female ventral black spot), we found strong enrichment for ESRRG-binding motifs and identified ASIP silencers containing ESRRG-binding sites (Fig. 5H). This is consistent with a model in which estrogen-responsive nuclear receptors repress ASIP in females, enabling local black pigmentation; whereas males lack such repression and retain uniform yellow ventral feather (Fig. 5I).

Motif enrichment also highlighted additional regulatory layers that may tune CRE coalition output (Fig. 5H): CLOCK implies rhythmic transcriptional inputs (*72–74*). MAX, an epigenetic sensor of cytosine modifications, suggests that the DNA modification state could contribute to this regulatory network (*75–77*). FOS::JUN and FOSL1::JUN are consistent with pioneer-factor activity (*78–81*). CTCF suggests local genome structure may contribute to CRE deployment (*81–85*). Together, results in Fig. 5 show that during complex patterning, ASIP expression is governed not by a single regulatory program but by flexible, modular interactions within CRE coalitions, whose accessibility varies across tissues, developmental stages, and sexes. Thus, CRE coalitions function as tissue-specific interpreters of upstream hormonal, rhythmic, epigenetic, and architectural inputs, enabling a single gene to generate diverse and dynamic pigment architectures—including stripes, spots, pinstripes with different size and spacing, or uniform coloration.

### 6. The AITE model: An Activator-Inhibitor Turing-Epigenetic (AITE) model integrates reaction-diffusion dynamics with epigenetic coalitions on a growing domain

The developmental, evolutionary, and cross-tissue analyses above converge on three interacting layers underlying ASIP patterning: (i) reaction-diffusion signaling, a Wnt-responsive transcriptional program, and secreted inhibitors; (ii) the expanding tissue geometry, which re-shapes inhibitory fields as the domain grows; and (iii) *cis*-regulatory logic, in which enhancer-silencer coalitions integrate multimodal signaling inputs into patterned transcription. What remains unclear is whether these components are individually necessary or jointly sufficient to generate the observed spatiotemporal sequence of stripes (A1→A2→A3) and their spacing on a growing tissue.

To test this, we sought a mathematical framework that unifies molecular signaling, tissue geometry, and *cis*-regulatory dynamics. We constructed an Activator-Inhibitor Turing-patterning Epigenetic (AITE) model that couples reaction-diffusion with CRE-coalition dynamics on a growing domain (Fig. 6A). The model includes five species 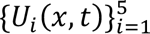 defined on a one-dimensional domain 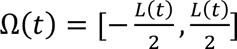, where each species evolves according to the reaction-diffusion equations:

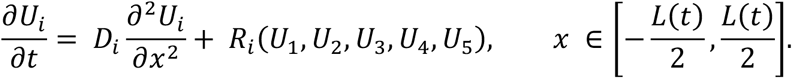

**Fig. 6.**
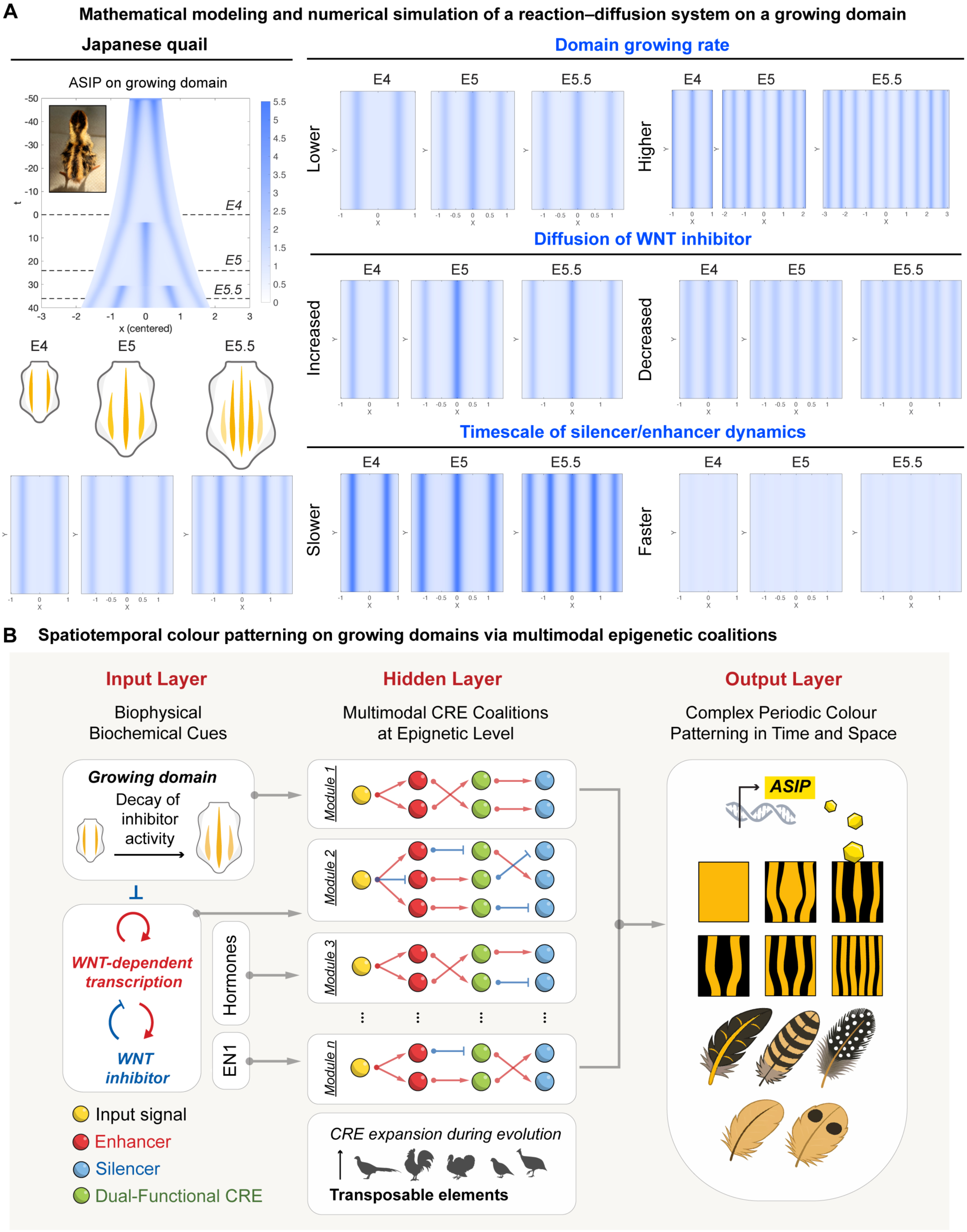
AITE model integrates reaction-diffusion dynamics with multimodal epigenetic CRE coalitions to generate periodic colour patterns. **A,** Mathematical modeling and numerical simulations of a reaction-diffusion system on a growing domain. Left: parameters used to model ASIP stripe formation in Japanese quail embryos. Right: altering parameters (e.g., domain growth rate, diffusion of Wnt inhibitor, or silencer timescale) changes the number, spacing, and intensity of ASIP stripes. **B,** Schematic illustration of the AITE (Activator-Inhibitor Turing-patterning Epigenetic) Model — a framework for periodic colour patterning through multimodal epigenetic coalitions of CREs, integrating tissue expansion and transposable-element inputs during evolution to generate complex colour patterns across time and space.

Here, the variables represent: WNT signaling (*U*_1_), WNT inhibitor (*U*_2_), ASIP expression (*U*_3_), ASIP enhancer activity (*U*_4_), and ASIP silencer activity (*U*_5_); *D_i_* is the diffusion coefficient of species *i*, and *R_i_* represents production, degradation, or interaction terms. Reaction terms describe production, degradation, or coupling among species, with enhancer and silencer terms modulating ASIP (U₃) through competitive interactions.

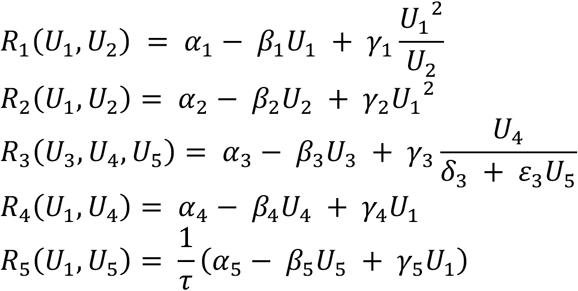

The domain length (tissue size) evolves in space in the form: 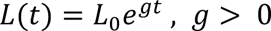

Parameter sweeps revealed intuitive control knobs in the integrated molecular system. As shown in Fig. 6A, in “Domain growing rate” panel, domain growth rate sets stripe timing and number: faster growth advances permissive thresholds and increases stripe count, whereas slower growth delays or reduces it. In “Diffusion of WNT inhibitor” panel, inhibitor diffusivity (Wnt antagonist spread) governs spacing: increased spread delays the appearance of new stripes and widens intervals; decreased spread accelerates emergence and narrows spacing. In “Timescale of silencer/enhancer dynamics” panel, CRE-coalition engagement timescale controls amplitude and persistence: slower silencer recruitment yields higher ASIP peaks and longer-lived stripes, whereas faster recruitment dampens peaks and can extinguish stripes. Critically, coupling growth-driven inhibitor dilution with enhancer-silencer dynamics recapitulates the sequential A1→A2→A3 progression on an expanding field, consistent with the delayed opening of silencers and the acquisition of ASIP-anchored chromatin loops observed from E4 to E5.5 (Fig. 1, E to H, and Fig. 6A).

Together, these analyses crystallize the AITE framework into four linked components (Fig. 6B): (i) Wnt lies upstream of ASIP transcription—epidermal ligands drive mesenchymal responses, ASIP-positive cells show elevated Wnt reporters; (ii) Stripe succession is growth-coupled—domain expansion dilutes inhibitors to open new permissive corridors (A1 → A2 at the midline → A3 between A1 and A2); (iii) CRE coalitions implement Turing-like logic—enhancers, silencers, and dual-function elements act on accessible chromatin whose positive and negative couplings convert graded signals into periodic transcription; (iv) patterning context dictates which CRE modules are engaged and how they assemble into CRE coalitions across tissues and stages.

More broadly, these analyses raise the possibility that the ASIP *cis*-regulatory landscape functions as a multilayer integrator—conceptually similar to hidden layers in a neural network—capable of combining diverse signaling inputs, growth dynamics, and chromatin constraints into discrete spatial outputs (Fig. 6B) (*86–88*). The widespread co-accessibility among enhancers and silencers, together with the mixture of positive and negative correlations across CREs, suggests an epigenetic architecture that could, in principle, support reaction-diffusion-like behavior at the level of chromatin regulation. While the AITE model is necessarily simplified, its ability to reproduce key qualitative features of stripe initiation, spacing, and succession provides a mechanistic basis for how classical patterning dynamics interface with *cis*-regulatory logic during development.

## Discussion

Colour patterns shape how organisms navigate their surroundings, respond to the physical world, and communicate within and between species. Pigment motifs act as ecological solutions: they may conceal, as in cryptic camouflage, or advertise, as in aposematic warning signals, and their evolution reflects predator behaviour, community context, and the physical environment (*89*). Environmental cues can also modulate these patterns, influencing developmental pathways and altering the physiological performance associated with pigmentation in birds (*90*, *91*). ASIP expression is highly sensitive to regulatory perturbation and strikingly dynamic, showing region-specific activity and rapidly shifting, periodically spaced patterns in developing avian skin. These features position ASIP as an evolutionary hotspot and a powerful paradigm for dissecting the epigenetic regulation of pigment patterning.

### 1. Avian skin as a platform for multi-scale colour patterning

Birds offer an exceptional system for dissecting ASIP regulation, as pigment patterns are organized hierarchically—from broad body domains to fine within-feather motifs. This multi-scale architecture provides a natural framework for linking distinct *cis*-regulatory modules to specific patterning outcomes. Recent single-nucleus multiome technologies, which jointly profile chromatin accessibility and gene expression, now enable regulatory elements to be resolved with single-cell precision and have already proven effective in diverse species and tissues (*92–96*).

In addition, avian pigment patterns form on growing domains—the expanding embryonic skin surface and the elongating filament in feather follicle—placing them in a regime where growth directly interacts with pattern formation. In such contexts, reaction-diffusion theory predicts shifts in instability modes, changes in wavelength selection, and characteristic transitions between pattern states as the domain expands (*10*, *97*, *98*). Together, these features position avian skin as a powerful platform for integrating biochemical and biophysical cues to reveal how the regulatory architecture of ASIP encodes complex pigment patterns.

### 2. Epigenetic coalitions and periodic ASIP expression

Here we reveal that coalitions of CREs allow ASIP to generate unique yet periodic and dynamic expression patterns. We define CRE coalition as a group of *cis*-regulatory elements—including enhancers, silencers, and dual-function elements—that act together at the epigenetic level to shape a specific pigment pattern. These coalitions operate on growing domains, from the expanding embryonic skin to the elongating feather filament, and their composition and activity change over developmental time. In the ASIP system, extensive co-accessibility among enhancers and silencers, linked through positive and negative couplings, establishes a chromatin-level regulatory syntax that, when embedded in a growing tissue, drives the sequential formation of ASIP stripes.

### 3. Regulatory grammar among CREs

Inspired by co-occurrence network analyses in microbial ecology (*71*), we applied analogous network-based reasoning to the ASIP *cis*-regulatory repertoire. Microbial studies show that non-random co-occurrence patterns can reveal potential positive and negative interactions within communities and illuminate the principles that govern their assembly (*70*). By analogy, we asked whether ASIP enhancers and silencers display reproducible co-activity patterns and what such associations might suggest about their regulatory relationships. It is well established that super-enhancers and super-silencers can form hubs with promoters to influence transcription (*27*, *32*, *40*, *99*, *100*). Building on these pioneering frameworks, we sought to explore additional questions: What is the communication logic within such epigenetic CRE coalitions? How does silencer-mediated repression differ from the functional shutdown of an enhancer? And how might these mechanisms operate in combination? Such a mixture of relationships among CREs may provide flexibility in how regulatory inputs are integrated during pattern formation without presupposing a single underlying mechanism. Our co-occurrence analysis offers only a preliminary view, and substantial experimental validation will be required to assess these possibilities.

As a conceptual aid, we draw an analogy to neural networks: biochemical and biophysical cues act as an input layer, feeding into a hidden layer of multimodal epigenetic CRE coalitions, and culminating in an output layer of pigment patterns (*86–88*). In a “bowtie architecture” view (*101*), diverse biochemical and biophysical inputs are compressed into a low-dimensional regulatory “waist” formed by CRE modules and then re-expanded into many pigment outputs, placing ASIP at the hub of a pattern-processing architecture. This framing also invites comparison to neural systems, where theories such as Neural Darwinism may offer inspiration for future work (*102*). Meanwhile, ecological metaphors such as the Red Queen hypothesis of co-evolving partners and the Black Queen hypothesis of functional dependency (*103*, *104*), providing useful parallels for thinking about CRE cooperation and competition. For example, elements brought into proximity by chromatin looping or shared transcription-factor usage may cooperate, whereas those that compete for factors or respond to opposing signals may act antagonistically.

### 4. Environmental modulation and CRE coalitions

New CRE coalitions may enhance the adaptability of ASIP expression to environmental and physiological cues, including sex hormones. Across vertebrates, colour frequently serves as a flexible signal of social standing. In fish such as cichlids, dominant males intensify their pigmentation, and changes in rank elicit rapid, hormone-driven colour shifts that broadcast competitive strength (*105*). Primates, as highly social species, use a similar strategy: in mandrills, the deep red facial and sexual skin of high-status males brightens as testosterone rises and fades when the alpha rank is lost. In both systems, ornamentation becomes a living readout of social position and underlying physiological state (*106*, *107*).

By analogy, hormone-responsive CRE coalitions at the ASIP locus could allow colour patterns to be tuned by systemic signals, coupling ecological and social context to local regulatory logic. Among pigmentation-related genes, ASIP stands out: nearly half of reported transposable-element-induced colour variants in mammals trace back to this locus (*17*), underscoring its sensitivity to regulatory innovation and its potential as a substrate for evolvable CRE coalitions.

### 5. Limitations and future directions

Based on these advances, there are important questions we can ask. How ASIP expression responds quantitatively to tissue growth, and whether the expanding domain itself acts as a biophysical cue that guides pigment patterning, remains incompletely understood. More broadly, it is not yet clear how tissues and developmental stages sculpt chromatin architecture to recruit distinct CRE coalitions, nor how these coalitions are reshaped by environmental and hormonal inputs *in vivo*. Addressing these limitations will require systematic perturbation of individual CREs and their combinations, live imaging of pattern dynamics in growing domains, and integrative modeling that links reaction-diffusion systems, chromatin topology, and CRE coalition logic. We aspire to understand the logic of pigment pattern formation on animal integument, and ASIP provides these new understanding at the epigenetic level.

## Acknowledgments

We thank all members of the Chuong laboratory for helpful discussions and support. During the preparation of this work the authors used ChatGPT to improve readability and language, using the following prompts: “polish the paragraph”. We do not use Generative AI to generate content. After using ChatGPT, the authors reviewed and edited the content as needed and take full responsibility for the content of the publication.

## Author contributions

Z.Y. and C.M.C. conceived the project, designed the experiments and wrote the manuscript. Z.Y. performed, analyzed, and interpreted most experiments. Cell culture, *in vitro* and *in vivo* reporter assays, somite injections, electroporation experiments, co-occurrence analysis were performed by Z.Y. W.Z. performed mathematical modeling with intellectual input from Q.N. Z.Y. and H.I.H. carried out single-nuclei Multiome experiment of E4 and E5.5 Japanese quail. Z.Y. and T.X.J. carried out single-nuclei Multiome experiment of adult guinea fowl and whole-mount *in situ*. Single-nuclei Multiome data were analyzed by C.K.C., W.Z., and Z.Y. Micro-C and bulk ATAC-seq were performed and analyzed by Z.Y., Y.C.L, and W.C.J. Transposable element and CRE conservation analyses across species were conducted by C.K.C. and Z.Y. Plasmid construction was designed by Z.Y. and carried out by Z.Y. with assistance from W.C.J., N.K.B, T.Y. Liu, and T.Y. Law. P.W. and T.Y. Liu provided assistance with Japanese quail husbandry. M.I. provided plasmids, performed whole-mount *in situ* hybridization, and provided intellectual input. Q.N. and E.C. provided intellectual input. All authors reviewed the manuscript.

## Data and materials availability

Further information and requests for resources and reagents should be directed to and will be fulfilled by the corresponding author, Dr. Cheng-Ming Chuong. Unique materials generated in this study are available from the corresponding author with a completed Materials Transfer Agreement. Any additional information required to reanalyze the data reported in this paper is available from the corresponding author upon request.

## Materials and Methods

### Ethics statement

All the animals used in this study were raised and handled according to authorized protocols (20231 and 21499) of the Institutional Animal Care and Use Committees of the University of Southern California (USC, Los Angeles, CA).

### Bird eggs

Japanese quail eggs were from AA laboratories (CA, USA). Guinea fowl eggs were from Fifth Day Farm, Inc. (PA, USA). Light Brown Leghorn chicken eggs were from Meyer Hatchery. All eggs were incubated at 38 °C and in 65% relative humidity until the specific stages.

### Sample preparation for snMultiome

For E4 and E5.5 Japanese quail, the embryonic dorsal skins were dissected, dissociated in 1 mL of dissociation solution containing 0.1% collagenase Type I and 0.25% Trypsin in 1x PBS and incubated in 37°C MultiTherm Shaker (Southern Labware), pipette every 5 min till there is no visible tissue. The dissociated cells were then filtered by a 70 μm and then a 40 μm cells trainers. 2x volume of chicken cell culture medium (DMEM with 10% FBS, 2% chicken serum, and 1% P/S) was added to eliminate the Trypsin reaction and cells were pelleted down by centrifugation at 300 x g for 10 min at 4°C. For adult guinea fowl feather, the feather tissue was dissociated in the same dissociation solution for 20 min, pipette every 5 min. Cells were washed in 1x cold PBS containing 2% BSA twice and cell numbers were count. Nuclei were isolated following the manufacturer’s instructions (10x Genomics, protocol CG000365 and CG000038, low cell input protocol), washed in wash buffer and resuspended in nuclei buffer. For each of samples, we generated separate barcoded single-nucleus libraries with simultaneous profiling of gene expression and chromatin accessibility using 10x Genomics Chromium Single Nuclei Multiome Reagent Kit, following the manufacturer’s instructions. Nuclei counting, suspension, single-nucleus libraries generation, and quality control were performed at the Molecular Genomics Core facility at University of Southern California Norris Comprehensive Cancer Center. The libraries were shipped to Novogene Corporation (CA, USA) and sequenced on the Illumina NovaSeq 6000 platform with paired-end 150 cycles.

### Multiomics analysis

Cellranger Arc v.2.0.0 (10x Genomics) was used for alignment and peak calling to generate a peak-by-cell and gene-by-cell count matrix. Reads were aligned to the Coturnix japonica 2.0 genome for Japanese quail, and NumMel1.0 for genome guinea fowl. Preprocessed digital matrices from the datasets were processed using Seurat (version 4.0.1) and Signac (version 1.12.0) (*93*). SoupX was used to remove the ambient RNA contamination while Doublet Finder was used to detect doublet (*108*, *109*). Seurat objects were generated and log-normalized using a scale factor of 10,000, and the top 2,000 variable features were identified with the vst method. Weighted Nearest Neighbor analysis was conducted to integrate multimodal RNA and ATAC datasets within individual single cells (*93*). DIRECT-NET, a machine-learning method based on gradient boosting, infers CREs by modeling the expression of a target gene as a non-linear function of the accessibilities of all candidate peaks (*53*). DIRECT-NET analysis was performed for each ASIP-positive mesenchymal population, including peaks located within ±250 kb of the ASIP transcription start site. For the DIRECT-NET-identified peaks, Pearson correlation coefficients with ASIP expression were calculated to classify each peak as an enhancer (positive correlation) or a silencer (negative correlation). CellChat was applied to assess the interaction strength among different cell types, with all analysis parameters kept at their default settings (*56*).

### Micro-C sample preparation

A Dovetail Micro-C Kit (Dovetail, #21006) was used for all Micro-C experiments. The procedure followed the manufacturer’s user guide (version 1.0) with minor modifications as previously described (*110*). Briefly, dissociated cells were sequentially cross-linked with DSG and formaldehyde, washed, digested with micrococcal nuclease, and lysed. A small aliquot was subjected to cross-link reversal and DNA fragment quality assessment, while the remaining lysate proceeded to proximity ligation. Chromatin was captured on magnetic beads and underwent end polishing, bridge adaptor ligation, and overnight intra-aggregate ligation. After cross-link reversal, ligated DNA was purified using SPRI beads. Sequencing libraries were prepared by end repair, Illumina adaptor ligation, and indexed PCR, followed by bead-based size selection (350-1000 bp) and final quality assessment before sequencing. The libraries were shipped to Novogene Corporation (CA, USA) and sequenced on the Illumina NovaSeq 6000 platform with paired-end 150 cycles.

### Micro-C analysis

Micro-C data were preprocessed according to Dovetail Micro-C Data Processing Guide. Briefly, BWA-MEM was used to map the sequencing reads to Coturnix japonica 2.0 genome. Valid pairs from the mapped reads were identified by pairtools parse pipeline. Unmapped, low mapping quality (quality <40), and PCR duplicate read pairs were removed. To visualize Micro-C data, juicertools.jar and Cooler tools were used to generate 1-kb resolution hic and cool contact matrices, respectively. Cool files were visualized using the Galaxy platform (*111*), version 22.1. The snapshots of contact matrices shown in this study were produced by Galaxy. Loops were discovered by Mustache with indicated resolution and FDR smaller than 0.2 (*112*).

### ATAC-seq sample preparation

Two biological replicates of each group were used. All procedures were performed on ice. ATAC resuspension buffer (ARB) was composed of 3 mM MgCl2, 10 mM NaCl, and 10 mM Tris-HCl (pH 7.4). Dissociated cells (50,000) of indicated tissues were lysed in 50 μl of cold ATAC lysis buffer (0.1% NP-40, 0.1% Tween-20, and 0.01% digitonin freshly added in ARB) for 5 min. One milliliter of ARB (with 0.1% Tween-20) was added and mixed well by inverting the tube. Nuclei were pelleted by 2500 rpm at 4°C for 10 min, and supernatants were discarded. Tn5 transposase (2.5 μl) was added to tagment chromatin at 37°C for 30 min in 50 μl of 1× Tagment DNA buffer (Illumina, #FC-121-1030) with 0.1% Tween-20, 0.01% digitonin, and 0.33× PBS. The tagmented DNA was purified using a DNA Clean & Concentrator-5 kit (Zymo Research, #D4014). To prepare the library, the tagmented DNA fragments were amplified in a thermocycler for 5 cycles using NEBNext High-Fidelity 2x PCR Mater Mix (NEB, #M0541S) in a total volume of 50 μl. Five microliters of the PCR products was used in qPCR to determine the number of extra amplification cycles before saturation, corresponding to 25% of maximum fluorescent intensity. The remaining 45 μl of the PCR products was amplified using the previously calculated cycle numbers and purified using 1x AMPure XP beads (Beckman, #A63880). For sequencing, the libraries were submitted to Novogene Corporation (CA, USA) and quantitated and analyzed using Qubit and Agilent 2100 Bioanalyzer, respectively. The libraries were shipped to Novogene Corporation (CA, USA) and sequenced on the Illumina NovaSeq 6000 platform with paired-end 150 cycles.

### ATAC-seq analysis

The ATAC-seq analysis was conducted on Partek Flow platform (Partek Inc.). Default parameters were applied. Prealignment QA/QC algorithms were run first. BWA-MEM algorithm v.0.7.17 (Genome: Coturnix japonica 2.0) was adopted and then analyzed by post-alignment QA/QC. Aligned reads were further filtered to remove duplicate alignments (defined as the same start and same sequence).

### RNA isolation and quantitative real-time PCR (qPCR)

Total RNA was extracted using Quick-RNA Miniprep Kit (Zymo, #R1054) according to the manufacturer’s protocols. For qPCR analysis, cDNA was synthesized using All-in-One 5X RT MasterMix (Abm, #G592) following manufacturer’s instructions. Quantitative PCR was performed using Maxima SYBR Green/ROX qPCR Master Mix (2X) (Thermo Fisher Scientific, #K0222) on a QuantStudio 3 Real-Time PCR System (Applied Biosystems, #A28567). PCR conditions were as follows: initial denaturation at 95°C for 10 minutes, followed by 40 cycles of 95°C for 15 seconds, 60°C for 30 seconds, and 72°C for 30 seconds. The delta-delta CT method was used to calculate the fold gene expression, with GAPDH serving as the internal reference gene. Primer sequences for qPCR were listed in table S2.

### Whole-mount *in situ* hybridization (WISH)

Whole mount *in situ* hybridization was done as described before (*113*). Briefly, embryos were fixed with 4% paraformaldehyde in PBS over night at 4°C, and then serially dehydrated with methanol. Hybridization with probes were performed over night at 65°C. Color development was done with NBT/BCIP substrate (Promega). RNA probes used in WISH were generated by PCR reactions from a mixture of E4 and E5.5 Japanese quail skin cDNA template. The primers are listed in table S2. PCR products were cloned into the pDrive vector (QIAGEN) to generate templates for digoxigenin-labeled probes, which were synthesized using T7 or U6 RNA polymerase (Roche).

### Cell culture

For *in vitro* culture of primary cells from dorsal and ventral Japanese quail skin, E7 embryos were used for cell harvest. Embryonic skin tissues were dissected and dissociated in 1 mL of dissociation solution containing 0.1% Type I collagenase and 0.25% trypsin in 1× PBS. Samples were incubated at 37 °C in a MultiTherm Shaker (Southern Labware) and triturated every 5 minutes until no visible tissue remained. The resulting cell suspension was sequentially filtered through 70 μm and 40 μm cell strainers. To quench trypsin activity, a 2x volume of chicken cell culture medium (DMEM supplemented with 10% FBS, 2% chicken serum, and 1% penicillin-streptomycin) was added. Cells were pelleted by centrifugation at 300 x g for 10 minutes at 4 °C. Primary Japanese quail cells and DF1 cell lines were maintained in chicken cell culture medium and cultured at 37 °C in an incubator with 5% CO2 and 95% air. Transfection of plasmids into cultured cells was performed using Lipofectamine 3000 (Thermo Fisher Scientific, #L3000015), following the manufacturer’s instructions. Reporter analysis was performed in the Flow Cytometry Facility at the Eli and Edythe Broad CIRM Center for Regenerative Medicine and Stem Cell Research at the University of Southern California, using an Attune NxT Flow Cytometer (Thermo Fisher Scientific, #A24858).

### *In vivo* electroporation for plasmid delivery

Japanese quail eggs were incubated till E2 for electroporation. The egg was windowed and a portion of the chorio-allantoic membrane removed to expose the embryo. The electrodes were placed closely adjacent to the two sides of the embryo’s body. A DNA solution (Tol2 plasmid: T2TP = 3:1 molar ratio, total DNA concentration 8 μg/μL) was injected into the neural tube of E2 stage. The electric pulses were delivered from a BTX Electro Square Porator ECM 830 through genetrodes (BTX model 512 with an angled 5mm gold tip) of 50 V, 20 ms, three times with 980 ms intervals. After treatment, the egg was sealed with scotch tape, and incubated for another 4 to 8 days at 38 °C.

### Plasmid construction

All Tol2 plasmids used here were generated by modifying pT2AL200R175-CAGGS-mCherry, pT2AL200R175-CAGGS-EGFP, and pT2AL200R175-CAGGS-H2BmCerulean provided by Dr. Masafumi Inaba (*16*). The reporter plasmids were assembled with the following step:

(1) miniCMV-d2GFP-bGH poly(A) fragment was PCR-amplified from plasmid 5HRE/GFP (Addgene, #46926). Chicken β-globin 5’HS4 insulator was PCR-amplified from plasmid GBSGFP insulator (Addgene, #121975). The two fragments are inserted into pT2AL200R175-CAGGS-mCherry digested with AflII and NsiI with In-Fusion Snap Assembly Master Mix (Takara, #638948) to assemble pT2AL200R175-miniCMV-d2GFP-insulator-CAGGS-mCherry.
(2) Chicken β-globin 5’HS4 insulator fragment 2 was PCR-amplified from plasmid GBSGFP insulator (Addgene, #121975) and inserted into pT2AL200R175-miniCMV-d2GFP-insulator-CAGGS-mCherry digested with MfeI and HpaI with In-Fusion Snap Assembly Master Mix (Takara, #638948) to assemble pT2AL200R175-miniCMV-d2GFP-insulator-CAGGS-mCherry-insulator.
(3) Chicken β-globin 5’HS4 insulator fragment 3 was PCR-amplified from plasmid GBSGFP insulator (Addgene, #121975) and inserted into pT2AL200R175-miniCMV-d2GFP-insulator-CAGGS-mCherry-insulator digested with AgeI and NsiI with In-Fusion Snap Assembly Master Mix (Takara, #638948) to assemble pT2AL200R175-insulator-miniCMV-d2GFP-insulator-CAGGS-mCherry-insulator.
(4) ASIP CRE fragments were PCR-amplified from E5.5 Japanese quail genomic DNA, inserted into the reporter backbone pT2AL200R175-insulator-miniCMV-d2GFP-insulator-CAGGS-mCherry-insulator digested with AgeI and NsiI with In-Fusion Snap Assembly Master Mix (Takara, #638948) between Chicken β-globin 5’HS4 insulator and miniCMV promoter.
(5) The insolation test plasmid, pT2AL200R175-insulator-d2GFP-insulator-CAGGS-mCherry-insulator, was assembled by inserting PCR-amplified fragment d2GFP-bGH poly(A)-insulator into pT2AL200R175-miniCMV-d2GFP-insulator-CAGGS-mCherry-insulator digested with SpeI and NsiI with In-Fusion Snap Assembly Master Mix (Takara, #638948), where miniCMV-d2GFP-bGH poly(A)-insulator was removed by digestion.

To generate Tol2-CAG-Jun-T2A-mCerulean, Tol2-CAG-EN1-T2A-mCerulean, and Tol2-CAG-EN1-T2A-EGFP, Jun and EN1 coding sequences were PCR-amplified from cDNA prepared from E4 and E5.5 Japanese quail embryos. T2A-mCerulean and T2A-EGFP fragments were amplified from pT2AL200R175-CAGGS-H2BmCerulean and pT2AL200R175-CAGGS-EGFP, respectively. Each construct was assembled by inserting the corresponding CDS together with the appropriate T2A fluorescent reporter into XhoI/ClaI-digested pT2AL200R175-CAGGS-H2BmCerulean with In-Fusion Snap Assembly Master Mix (Takara, #638948).

### Correlation-based co-occurrence CRE network analysis

#### (1) ATAC-seq Signal Extraction

ATAC-seq signal intensity across predefined genomic regions (CRE regions) were quantified. First, BED and bedGraph files were coordinate-sorted. Sorted bedGraph files were then converted into bigWig format using bedGraphToBigWig (*114*) with the appropriate genome size file. Per-region signal metrics (including length-weighted mean signal) were computed for each bigWig file using bigWigAverageOverBed (*114*). The resulting values were merged into a single matrix, preserving the original region order. This matrix provides mean ATAC-seq signal per sample for every genomic region and was used for downstream analyses.

#### (2) Co-occurrence Network Generation

A correlation-based co-occurrence analysis was performed to identify CREs exhibiting coordinated patterns of chromatin accessibility across samples. The analytical principles underlying this approach have been described previously (*115*). ATAC-seq signal values (normalized accessibility per CRE per sample) were assembled into a matrix with rows representing CREs and columns representing biological samples. To reduce noise from extremely low-accessibility elements, CREs were first ranked by their total accessibility (row-sum) across all samples. The top 80% of CREs were retained for downstream analysis, ensuring that the correlation structure was driven by CREs with reliable and biologically meaningful ATAC-seq signal. The resulting matrix was transposed and rank-normalized, generating a Spearman-like representation suitable for non-normally distributed accessibility data. Pairwise correlations between all CREs were computed, and for each CRE pair a correlation coefficient (r), t-statistic, and p-value were calculated. P-values were subsequently corrected for multiple testing using the Benjamini-Hochberg FDR method. Only CRE pairs exhibiting both strong correlations (|r| ≥ 0.5) and statistical significance (FDR < 0.01) were retained. The thresholded correlation matrix was converted into an undirected igraph network in which nodes correspond to CREs and edges denote significant co-accessibility (positive r) or mutually exclusive accessibility (negative r). Signed correlations were retained as edge attributes, while absolute correlation values were used as weights. CRE metadata (e.g., functional annotations) were added to each node. The final edge and node tables were exported for visualization and downstream network analysis.

#### (3) Co-occurrence Network Visualization

Co-occurrence networks were visualized using Gephi (version 0.10.1) (*116*). Network layouts were generated with the Fruchterman-Reingold force-directed algorithm, which positions nodes in two-dimensional space based on balanced attractive and repulsive forces to produce an interpretable representation of an undirected graph (*117*). Nodes and edges were color-coded according to CRE metadata (e.g., functional annotations, correlations, ATAC-seq signal). Node size was scaled in proportion to each node’s degree to highlight highly connected CREs.

#### (4) Weighted Network Community Detection (Louvain Modularity)

We conducted community detection using the Louvain modularity optimization algorithm in Gephi (version 0.10.1) (*116*, *118*). The analysis was performed with edge weights enabled and randomized processing order to avoid local minima. The modularity resolution parameter was set to 1.0, corresponding to the standard resolution used in the Louvain method. The algorithm returned a modularity value of 0.440, revealing a moderately modular network structure. In total, four communities were detected, each representing clusters of nodes with higher internal than external connectivity.

#### (5) Detection of Sample-Enriched CRE Modules

To identify sample-enriched accessibility signatures among CRE modules, a matrix of average ATAC-seq signal values for each modularity class across all samples was evaluated. For each modularity class, accessibility values across samples were standardized by converting them to row-wise z-scores, allowing comparison of relative enrichment patterns independently of overall signal magnitude. Modules showing selectively elevated accessibility in a single sample relative to all others were classified as sample-specific signatures, indicating distinct regulatory activity in that condition. Both the set of exclusive sample-enriched modules and the full collection of enriched module-sample pairs were compiled for downstream interpretation. This analytical framework enabled systematic identification of CRE modules exhibiting preferential accessibility in specific developmental stages or tissue contexts.

### Mathematical modeling and simulation

To facilitate the numerical simulations, we introduce the rescaled spatial coordinate 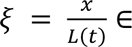 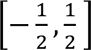 and define 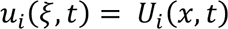 which maps the system from the expanding physical domain to a fixed reference domain, yielding the transformed reaction-diffusion equations:

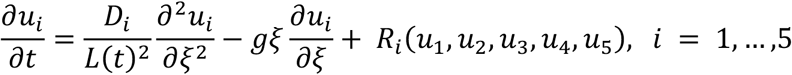

This formulation highlights the combined effects of diffusion (with effective diffusivity 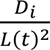), advection due to exponential growth 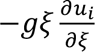, and the reaction kinetics *R_i_*.

The full parameters used in modeling and their descriptions are listed in table S3. The growth rate is estimated from the quail embryo lengths at E4, E5, E5.5 (*16*), using a least-squared fitting approach. The remaining parameters were chosen to reproduce the desired spatial patterns.

For numerical simulations, the reaction-diffusion system for the five-species model was discretized in space using a uniform grid with N=100 points on the fixed reference domain ζ ∈ 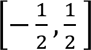. Second-order finite differences were used to approximate the spatial derivatives, and zero-flux (i.e., Neumann) boundary conditions were enforced via mirrored ghost points. The resulting large system of coupled ordinary differential equations in time was solved using MATLAB’s ODE15s solver, which efficiently handles stiff dynamics arising from disparate diffusion rates and nonlinear reaction kinetics. Simulations were initialized with small random perturbations around the homogeneous steady state to trigger pattern formation.

### Statistical analyses

Statistical analyses were performed using GraphPad Prism v.8.4.2 (GraphPad Software). An unpaired two-sided Student’s t-test was used to compare groups. Data are presented as mean ± standard deviation of the mean (s.d.). Statistical significance was designated as follows: *p < 0.05, **p < 0.01, ***p < 0.001, ****p < 0.0001.

