## Supplementary material for "Epigenetic Coalitions Couple Tissue Growth to Generate Periodic Colour Patterns in Birds": fig. S

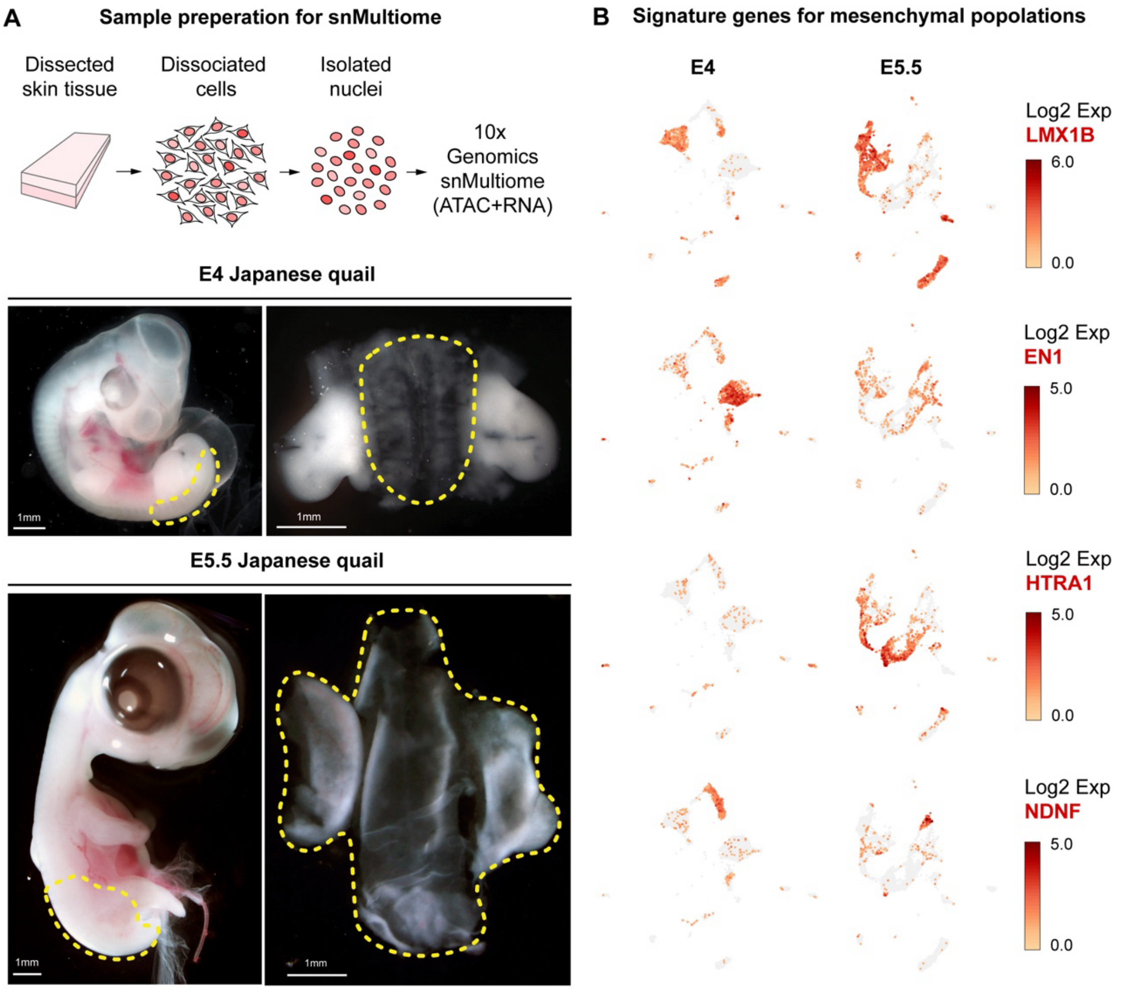


**Fig. S1. Tissue collection and mesenchyme population clustering.**

**A,** Skin samples from E4 and E5.5 embryos were collected following a standardized workflow. Skin tissue was dissected from the dorsal region, with the yellow dashed line indicating the precise domain isolated for following processing. Dissected tissue was enzymatically and mechanically dissociated to obtain a single-cell suspension, followed by gentle lysis to release intact nuclei. Purified nuclei were then processed for snMultiome profiling to jointly capture chromatin accessibility and gene expression at single-nucleus resolution.

**B,** Expression profiles of signature genes distinguish the four mesenchymal populations: ML is marked by LMX1B, M1 by EN1, M2 by NDNF, and M3 by HTRA1.



INTERNAL

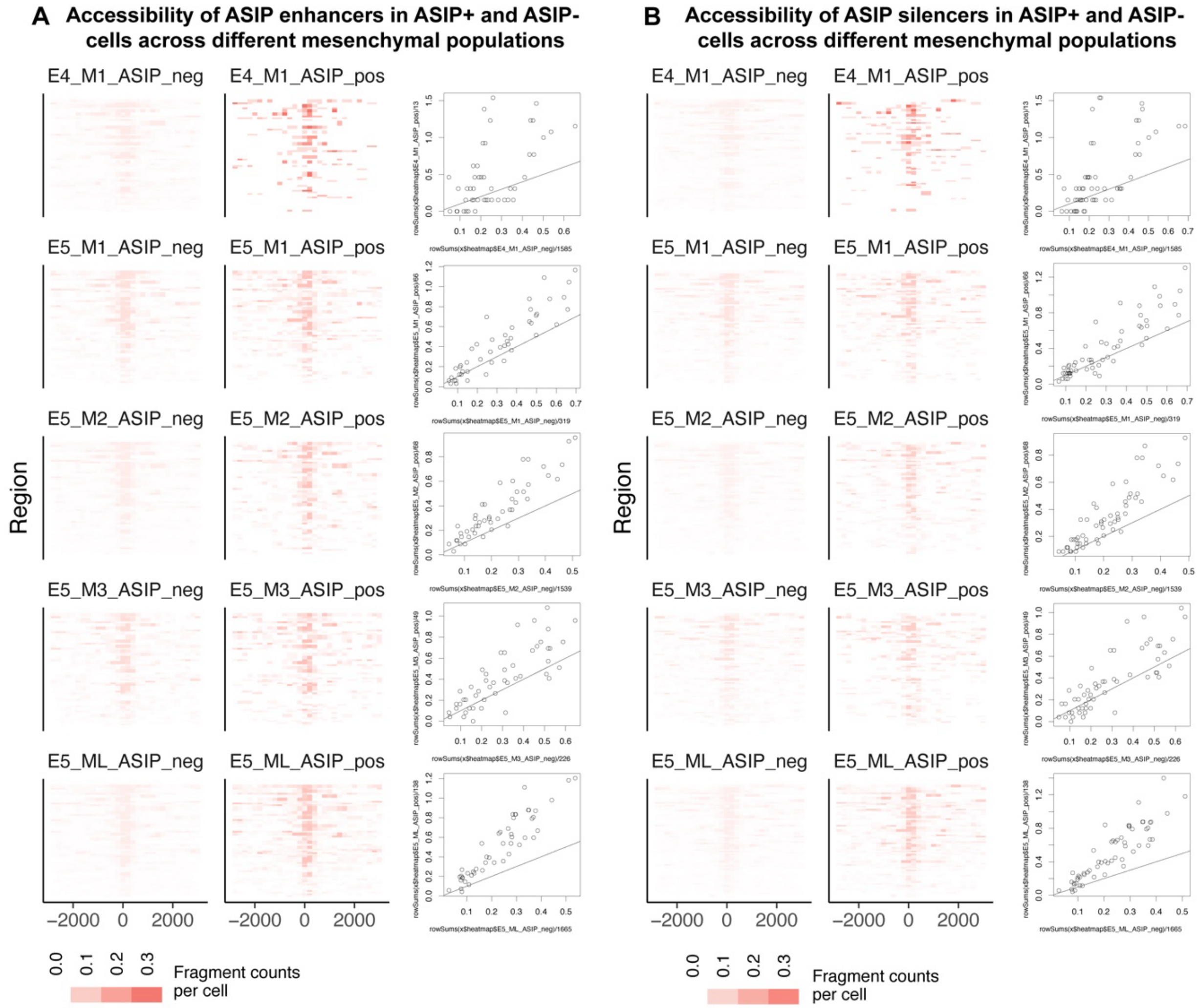


**Fig. S2. Accessibility of ASIP CREs in ASIP+ versus ASIP- cells across mesenchymal subtypes.**

**A,** Accessibility of ASIP enhancers in ASIP+ versus ASIP- cells across mesenchymal populations.

**B,** Accessibility of ASIP silencers in ASIP+ versus ASIP- cells across mesenchymal populations.



INTERNAL

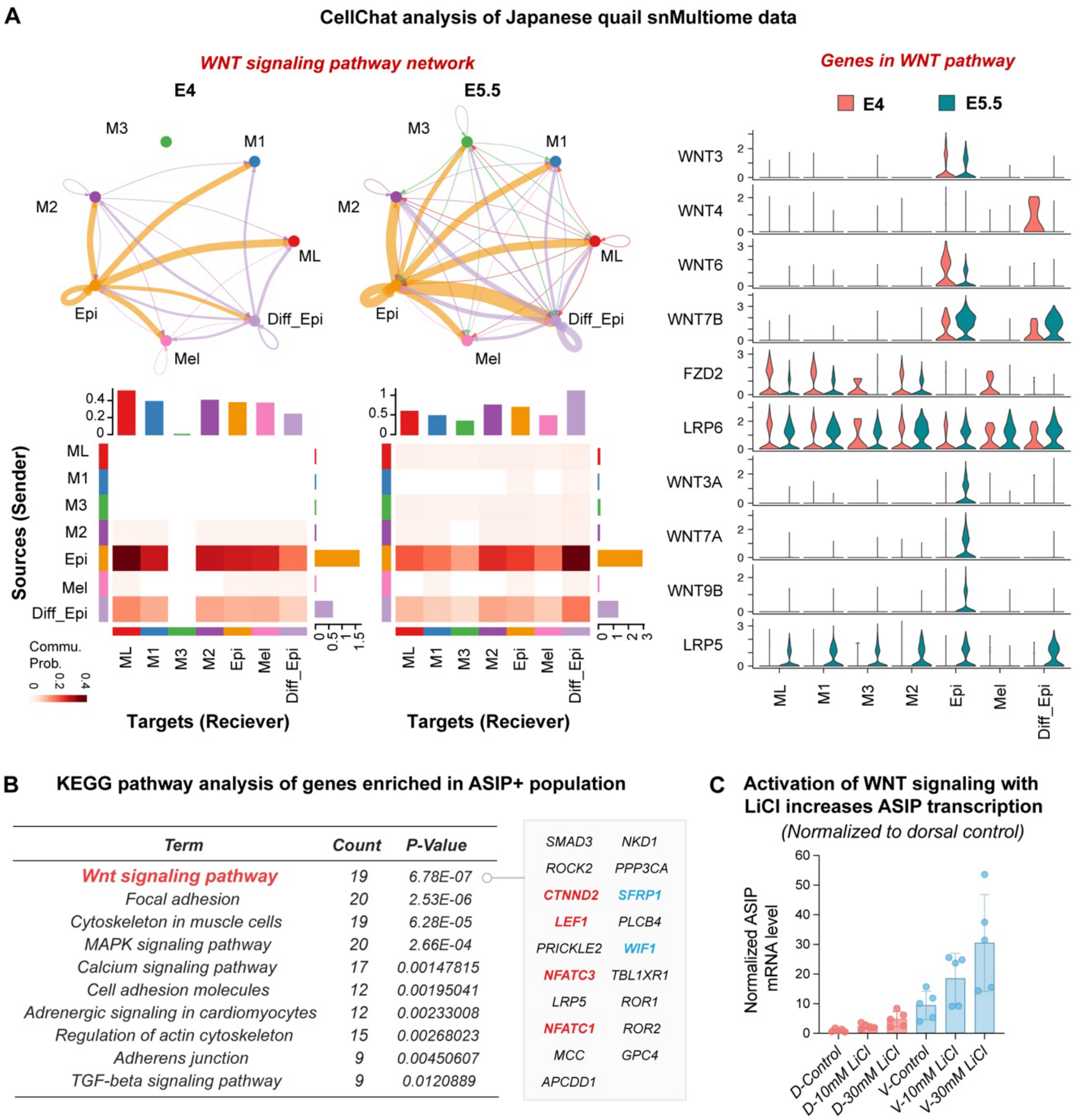


**Fig. S3. WNT signaling acts as an upstream regulator of ASIP expression.**

**A,** CellChat analysis of Japanese quail snMultiome data indicates that the epidermis acts as the source of Wnt ligands (WNT3, WNT4, WNT6, WNT7B, WNT3A, WNT7A, WNT9B), while the mesenchyme serves as the receiver, expressing the corresponding receptors and co-receptors (FZD2, LRP5, and LRP6).

**B,** KEGG pathway analysis of genes enriched in ASIP+ population.

**C,** Activation of WNT signaling with LiCl increases ASIP transcription (normalized to dorsal control).



INTERNAL

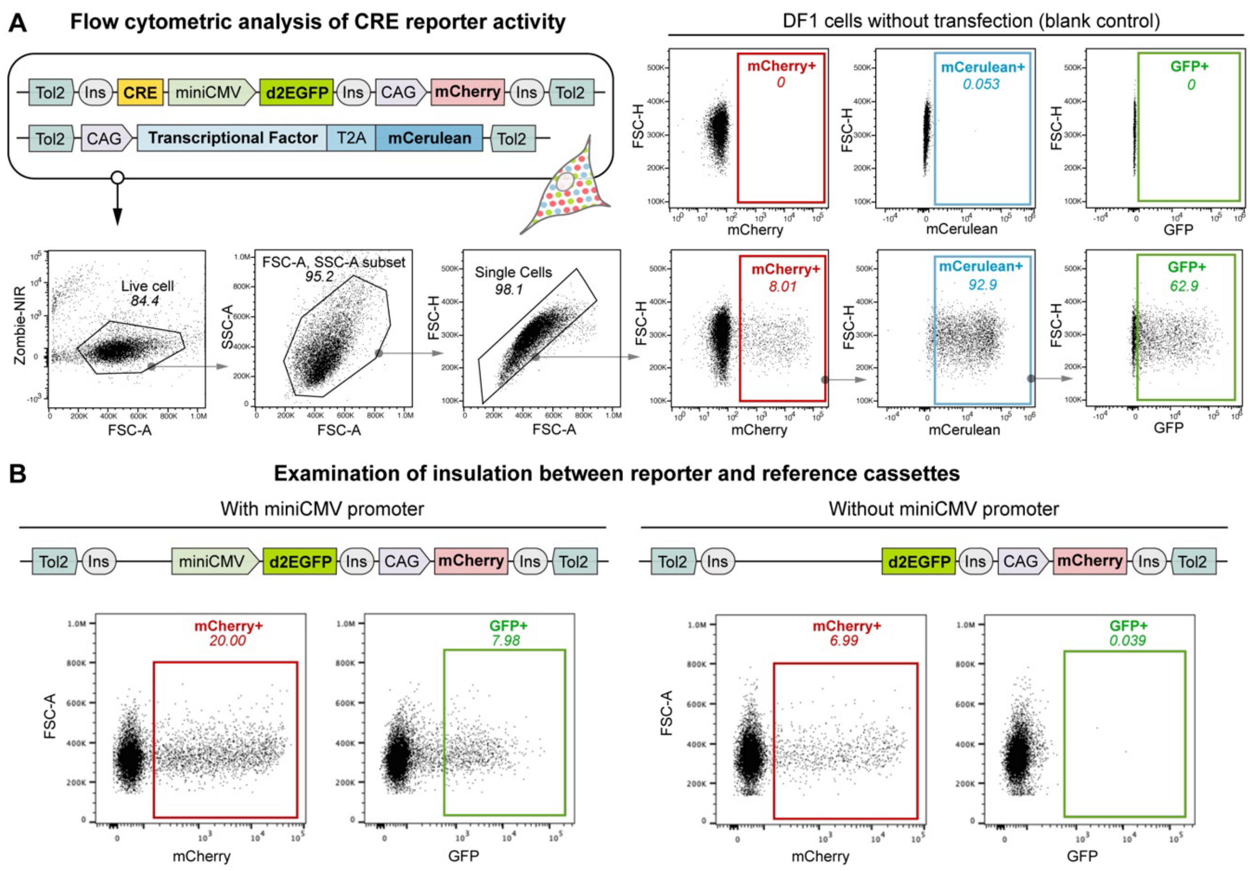


**Fig. S4. Functional characterization of CRE reporter activity and cassette insulation.**

**A,** Flow cytometric analysis of CRE reporter activity.

**B,** Examination of insulation between reporter and reference cassettes. Removing the miniCMV from the d2GFP reporter cassette abolished d2GFP expression, confirming that no signal arises from leakage of CAG promoter activity in the mCherry reference cassette. These results indicate that the HS5′ insulator provides robust insulation between the reporter and reference cassettes.



INTERNAL

A

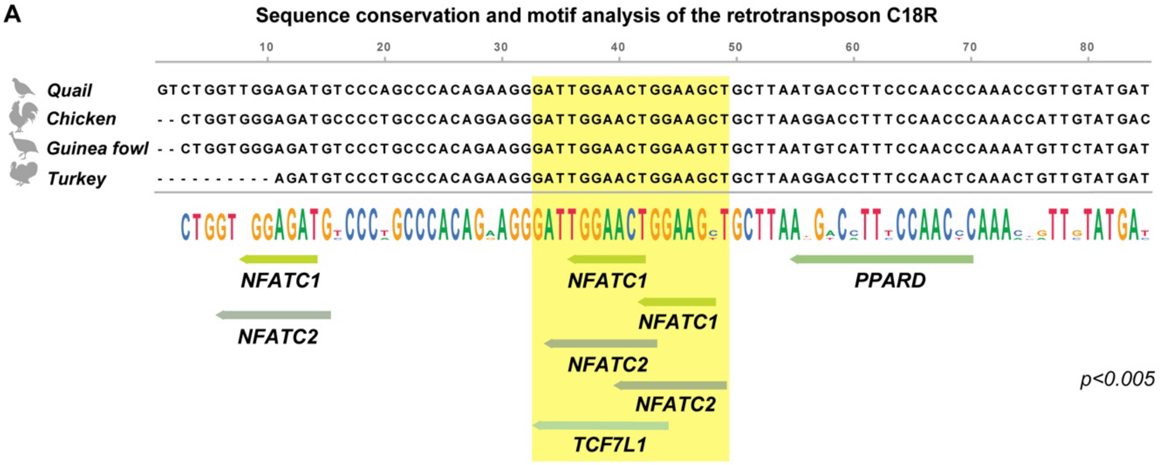


B

| Motif ID | Alt ID | Sequence Name | Strand | Start | End | p-value | q-value | Matched Sequence |
|---|---|---|---|---|---|---|---|---|
| MA0624.1 | NFATC1 | C18R | - | 34 | 43 | 0.00168 | 0.144 | CAGTTCCAAT |
| MA0624.1 | NFATC1 | C18R | - | 40 | 49 | 0.00195 | 0.144 | AGCTTCCAGT |
| MA0624.1 | NFATC1 | C18R | - | 6 | 15 | 0.00497 | 0.194 | CATCTCCAAC |
| MA0624.1 | NFATC1 | C18R | + | 59 | 68 | 0.00549 | 0.194 | CCTTCCCAAC |
| MA0624.1 | NFATC1 | C18R | + | 58 | 67 | 0.00775 | 0.194 | ACCTTCCCAA |
| MA0624.1 | NFATC1 | C18R | + | 13 | 22 | 0.00791 | 0.194 | ATGTCCCAGC |

| Motif ID | Alt ID | Sequence Name | Strand | Start | End | p-value | q-value | Matched Sequence |
|---|---|---|---|---|---|---|---|---|
| MA0152.1 | NFATC2 | C18R | - | 36 | 42 | 0.00158 | 0.128 | AGTTCCA |
| MA0152.1 | NFATC2 | C18R | - | 42 | 48 | 0.00169 | 0.128 | GCTTCCA |
| MA0152.1 | NFATC2 | C18R | - | 8 | 14 | 0.0025 | 0.128 | ATCTCCA |
| MA0152.1 | NFATC2 | C18R | + | 59 | 65 | 0.00549 | 0.211 | CCTTCCC |

| Motif ID | Alt ID | Sequence Name | Strand | Start | End | p-value | q-value | Matched Sequence |
|---|---|---|---|---|---|---|---|---|
| MA1421.1 | TCF7L1 | C18R | - | 33 | 44 | 0.00329 | 0.378 | CCAGTTCCAATC |
| MA1421.1 | TCF7L1 | C18R | - | 5 | 16 | 0.00524 | 0.378 | ACATCTCCAACC |
| MA1421.1 | TCF7L1 | C18R | + | 18 | 29 | 0.00955 | 0.459 | CCAGCCCACAGA |

| Motif ID | Alt ID | Sequence Name | Strand | Start | End | p-value | q-value | Matched Sequence |
|---|---|---|---|---|---|---|---|---|
| MA1550.1 | PPARD | C18R | - | 55 | 70 | 0.000454 | 0.058 | GGGTTGGGAAGGTCAT |
| MA1550.1 | PPARD | C18R | + | 4 | 19 | 0.00863 | 0.552 | TGGTTGGAGATGTCCC |

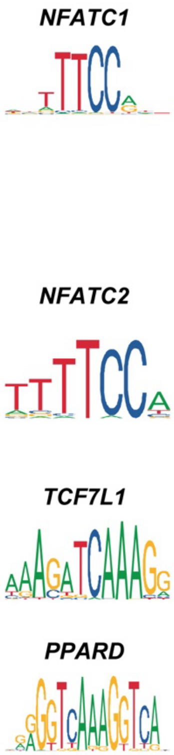


**Fig. S5. Sequence conservation and transcription-factor motif architecture of the retrotransposon C18R.**

**A,** Sequence conservation and motif analysis of the retrotransposon C18R.

**B,** Detailed motif analysis for NFATC1, NFATC2, TCF7L1, and PPARD within the retrotransposon C18R sequence ($p < 0.01$).



INTERNAL

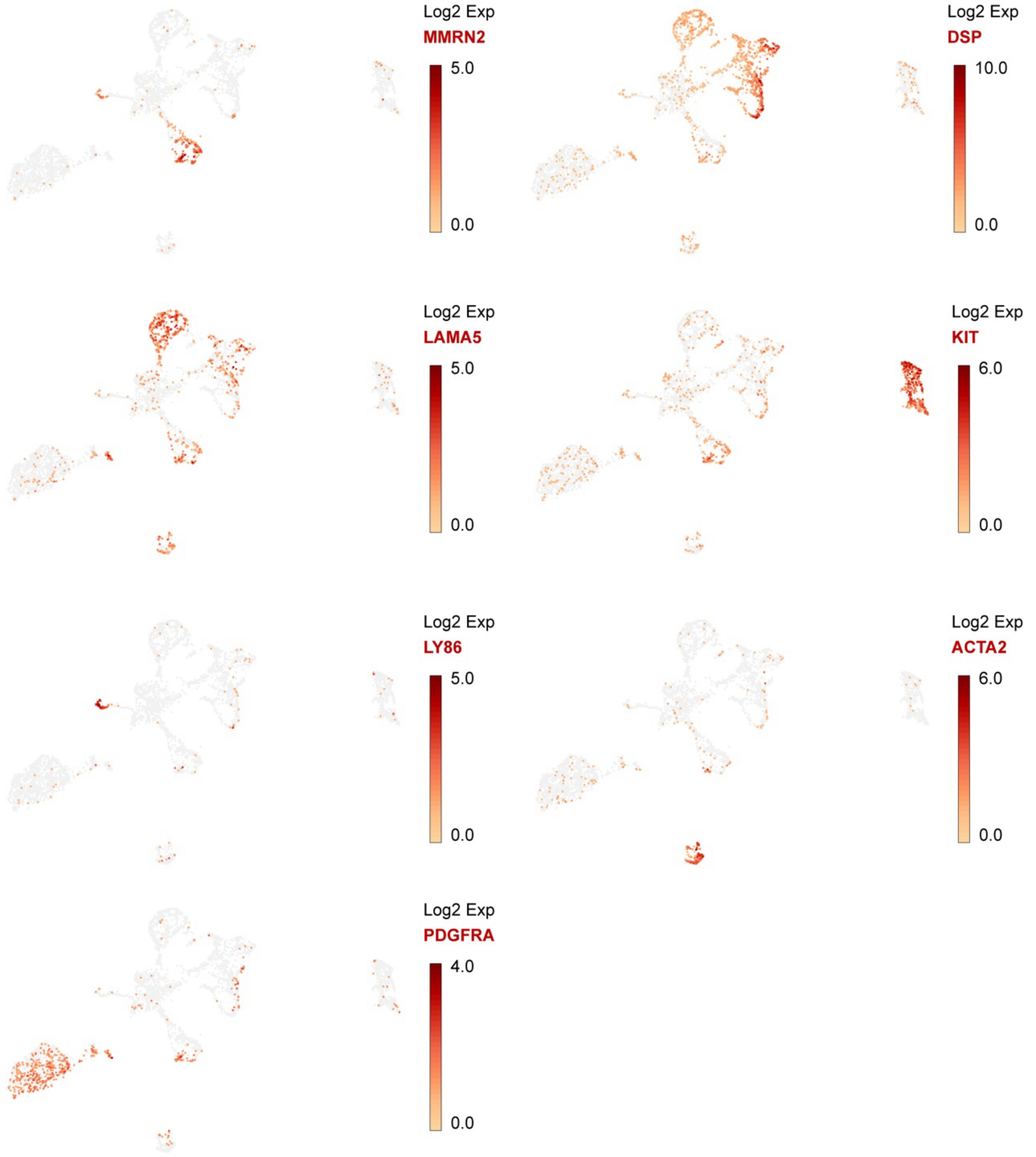


**Fig. S6. Expression profiles of signature genes distinguish cell clusters.** Endothelial cells are marked by MMRN2, epidermis 1 by DSP, epidermis2 by LAMA5, immune cells by LY86, melanocytes by KIT, mesenchymal cells by PDGFRA, and myocytes by ACTA2.



INTERNAL

**Table S1. 10 transposable elements identified within the Guinea fowl ASIP CREs**

| Chr | begin | end | name | class/family | id | CRE |
|---|---|---|---|---|---|---|
| NC_034427.1 | 1306842 | 1307199 | CR1-Y2_Aves | LINE/CR1 | 390823 | GF_CRE21 |
| NC_034427.1 | 1316690 | 1317090 | CR1-15_Crp | LINE/CR1 | 390829 | GF_CRE4 |
| NC_034427.1 | 1320998 | 1321409 | CR1-Y2_Aves | LINE/CR1 | 390833 | GF_CRE16 |
| NC_034427.1 | 1374538 | 1374778 | GGLTR5A | LTR/ERVL | 390850 | GF_CRE27 |
| NC_034427.1 | 1544057 | 1544536 | GGLTR5A | LTR/ERVL | 390924 | GF_CRE10 |
| NC_034427.1 | 1544544 | 1544906 | CR1-Y2_Aves | LINE/CR1 | 390925 | GF_CRE10 |
| NC_034427.1 | 1589889 | 1590137 | CR1-Y2_Aves | LINE/CR1 | 390947 | GF_CRE14 |
| NC_034427.1 | 1631502 | 1631645 | UCON2 | DNA? | 390959 | GF_CRE22 |
| NC_034427.1 | 1693077 | 1693436 | CR1-Y1_Aves | LINE/CR1 | 390986 | GF_CRE25 |
| NC_034427.1 | 1693830 | 1693926 | Chompy-4_Croc | DNA/PIF-Harbinger | 390987 | GF_CRE25 |



INTERNAL

**Table S2. Primer list with sequence (5’-3’)**

| | |
|---|---|
| **Primers for in situ probe cloning** | |
| LMX1B_JQ_Pb2_F | CCCCATATGGACATCGCCTCGG |
| LMX1B_JQ_Pb2_R | AATCAAGGGTATGTGACTGATGGATATGTTTTGT |
| EN1_JQ_Pb_1F | GCCTCAACGAGTCCCAGATC |
| EN1_JQ_Pb_1R | AAGGATCGAACGGAAGAGCG |
| HTRA1_JQ_Pb1_F | CATCAATTATGGAAACTCTGGAGGACCC |
| HTRA1_JQ_Pb1_R | TACTGTCCTTTTTAATAATGTCACTGACATCAGAGGC |
| NDNF_JQ_Pb2_F | GCCAGTGGAGAAGGTTCAGGT |
| NDNF_JQ_Pb2_R | TGTGTCGGGCTTCAGGTCAG |
| JQ-ASIP-AS-T7 | CTAATACGACTCACTATAGGGAGAGAAAAATTCGGCATTT GCAC |
| JQ-ASIP-S1 | TGAGGTTTTGACTGACCTTGG |
| **Primers for qPCR** | |
| GAPDH-JQ-F | GGAGAAACCAGCCAAATATGATG |
| GAPDH-JQ-R | AGGTGGAAGAATGGCTGTCA |
| qPCR-ASIP-F1 | ATCTCCCACCCATCTCCATC |
| qPCR-ASIP-R1 | TGGGGGTGTCTTCAGTTCAG |



INTERNAL

**Table S3. Parameters of the five-species reaction-diffusion model.**

| Parameter | Description | Value |
|---|---|---|
| $g$ | Domain growth rate (mm/hr) | 0.0155323 |
| $D_1$ | Diffusion coefficient of WNT signaling | 0.0013 |
| $D_2$ | Diffusion coefficient of WNT inhibitor | 0.0398 |
| $D_3$ | Diffusion coefficient of ASIP | 0.0066 |
| $D_4$ | Diffusion coefficient of ASIP enhancer | 8.8064e-06 |
| $D_5$ | Diffusion coefficient of ASIP silencer | 8.8064e-05 |
| $\alpha_1$ | Basal production rate of WNT signaling | 1 |
| $\beta_1$ | Decay rate of WNT signaling | 10 |
| $\gamma_1$ | Auto-activation strength of WNT signaling | 10 |
| $\alpha_2$ | Basal production rate of WNT inhibitor | 1 |
| $\beta_2$ | Decay rate of WNT inhibitor | 10 |
| $\gamma_2$ | Activation strength of WNT inhibitor by WNT signaling | 10 |
| $\alpha_3$ | Basal production rate of ASIP | 4 |
| $\beta_3$ | Decay rate of ASIP | 10 |
| $\gamma_3$ | Activation strength of ASIP by ASIP enhancer | 10 |
| $\delta_3$ | Basal threshold of ASIP inhibitor effect on ASIP production | 0.1 |
| $\varepsilon_3$ | Strength of ASIP silencer inhibiting ASIP | 1.35 |
| $\alpha_4$ | Basal production rate of ASIP enhancer | 1 |
| $\beta_4$ | Decay rate of ASIP enhancer | 10 |
| $\gamma_4$ | Activation of ASIP enhancer by WNT signaling | 10 |
| $\alpha_5$ | Basal production rate of ASIP silencer | 1 |
| $\beta_5$ | Decay rate of ASIP silencer | 10 |
| $\gamma_5$ | Activation of ASIP silencer by WNT signaling | 10 |
| $\tau$ | Time scale factor for ASIP silencer dynamics | 1000 |



INTERNAL
